# Effects of dead conspecifics, hunger states, and seasons on the foraging behavior of the purple urchin *Heliocidaris crassispina*

**DOI:** 10.1101/2020.03.09.984674

**Authors:** Dominic Franco C. Belleza, Yuuki Kawabata, Tatsuki Toda, Gregory N. Nishihara

## Abstract

Trophic cascades exerts a powerful effect between predator and prey relationships in an ecosystem. In aquatic environments, the signals associated with predators and predation are used by prey as a cue to avoid encountering predators when foraging for food. These cues are powerful enough to control prey populations and indirectly protect primary producers. We evaluated the effects of cues associated with predation on the purple urchin, *Heliocidaris* crassispina and examined effects of hunger state and season using time-lapse photography, we conducted a series of manipulative and *in situ* behavior experiments to determine foraging behavior patterns which demonstrate behavior modification. The results suggest that starved urchins were less sensitive to predation cues when compared to normally fed urchins. Field experiments indicated that 70% of fed urchins fled when exposed to a predation cue (presence of a dead urchin), whereas all starved urchins remained regardless of the cue, supporting the results from the laboratory using the dead urchin and algae treatment cues. Sea urchin activity and feeding rates were lower in winter-spring than in summer-autumn. We suggest that hunger state has a large influence over the behavioral-response of sea urchins, while also being affected by season due to metabolic control. In general, starvation overrides predator avoidance behaviors and exposes prey species to higher risks of predation.

## 1. INTRODUCTION

Predation and resource availability control food webs in the ecosystem (Nielsen & Navarrete 2004, Lynam et al. 2017). These interacting forces, together with variabilities in environmental stress depend on the regulating effect (i.e., energy allocation, expenditure, and transfer) they exert on the community of producers and consumers (Menge & Sutherland 1987). Hairston et al. (1960) hypothesized that populations of herbivores and the level of herbivory were generally controlled by predation rather than by food supply (i.e., “green world” hypothesis) and therefore the collapse of predator populations increased the likelihood of herbivore domination (Estes & Palmisano 1974).

In temperate regions, macroalgal forests are an important coastal ecosystem (Steneck et al. 2002, Smale et al. 2010, Langlois et al. 2011). Within the canopy, the high diversity of fish and invertebrate are dependent on the canopy for food and refuge (Lowry & Pearse 1973, Holbrook et al. 199, Kamimura & Shoji 2009). The sea urchin is a keystone species in marine forests, because they can overwhelm net benthic primary production (Tuya et al. 2004). In ecosystems where apex predator populations are intact, urchin populations are maintained through predation (Tegner & Levin 1983, Pearse & Hines 1987, Sala & Zabala 1996, Sievers & Nebelsick 2018). When predation pressure is removed, the urchin population leads to overgrazing, eventually converting seaweed beds into barrens. In general, barren areas are characterized by low diversity and habitat complexity (Mangialajo et al. 2008). Large canopy forming macroalgae are replaced by grazing resistant turf forming macroalgae (Wright et al. 2005), considered to be an intermediary stable state supported by strong feedback mechanisms (Filbee-Dexter and Wernberg 2018). As grazing pressure surpasses the thresholds of the remaining primary producers, the community state eventually transitions into a species-poor stable state (Steneck et al. 2002, Filbee-Dexter & Scheibling 2014; Ling et al., 2015) that is easily maintained by a few urchins (Tuya et al. 2004, Bonaviri et al. 2011). However, sustained human intervention or the recovery of predator populations can revert barrens into a macroalgal-dominated state (Blamey et al. 2013, Steneck et al. 2013).

Attempts to revert barrens into seaweed forests are not uncommon. Methods include manual removal or destruction of urchins and other herbivores to encourage natural recruitment of juvenile seaweeds (Yotsui & Maesako 1993, Watanuki et al. 2010, Nanri et al. 2011), small-to-medium scale transplantation of fertile seaweed thalli and mass dispersal of viable spores (Hernandez-Carmona et al. 2000, Yoon et al. 2013, Ogata et al. 2016), and installment of artificial reefs (Watanuki & Yamamoto 1990, Westermeier et al. 2013). Experimental evidence has shown that human intervention may succeed and promote seaweed forest recovery (Ling et al. 2010; Verdura et al., 2018; Layton et al., 2020; Verges et al., 2020). However, maintaining restored algal forests becomes difficult when uncontrolled urchin population levels eventually establish dense feeding fronts (Lauzon-Guay & Scheibling 2007, Ling & Johnson 2009). Regardless of the situation, the decision to restore ecosystems must be evidence-based and scale and context-specific (Johnson et al. 2016).

Harnessing the effect of natural predators on prey to indirectly maintain the population of primary producers may be a more practical solution (Schmitz et al. 2004). The direct reduction in the population of herbivores through consumption is called density-mediated indirect interaction (DMII) while the modification of prey behavior is called trait-mediated indirect interaction (TMII) (Schmitz et al. 2004). These interactions were observed in a variety of terrestrial and aquatic ecosystems (Shurin et al. 2002). In the aquatic ecosystem, the effects of trophic cascade seem to be more prominent than in terrestrial ecosystems (i.e., marine benthos > marine plankton > terrestrial food web) (Strong 1992, Halaj & Wise 2001, Shurin et al. 2002). The non-lethal effect of TMII may be comparable in magnitude to that of DMII, because behavior change has population-wide effects, whereas direct predation only affects the individual (Peacor & Werner 2001; Pessarrodona et al. 2019).

Historically, algal forests composed of large brown algae created dense expansive belts around the coastline of Japan and supported a large diversity of economically important fish and invertebrates (Uki et al. 1986, Kamimura & Shoji 2009). Presently, seaweed forests in Japan are undergoing a catastrophic decline (“isoyake”) and the remaining seaweed forests are at high risk (Okuda 2008, Haraguchi & Sekida 2008, Fujita 2010). The loss of seaweed forests has led to the decline in coastal fisheries production (Kiyomoto et al. 2013). Efforts to revert the decline in seaweed forests has produced numerous guidelines and methodologies, however success is limited (Terawaki et al. 2003, Fujita 2010, Kuwahara et al. 2010, Fujita 2015).

We focused on determining the impact of a non-lethal perceived threat on the foraging behavior of the purple urchin *Heliocidaris* crassispina Agassiz. Experimental studies have shown urchins to have complex foraging behaviors (Vanderklift & Kendrick 2005, Kriegisch et al 2019) and that negative responses from manipulative experiments ranged from strong (Campbell et al. 2001, Hagen et al. 2002) to weak (Harding & Scheibling 2015). Here we use dead conspecifics as a deterrent (Campbell et al 2001, Morishita & Barreto 2011) to explore the effects of the prey species’ hunger state and season on the urchin’s decision-making process. The individual’s hunger state (i.e. satiety vs. starvation) has been known to modulate an individual’s perception of risk (Clark 1994) while the season is associated with reproductive phenology (Agatsuma et al. 2000, Yatsuya & Nakahara 2004a). In this study we examine prey behavior towards predation signals in better understanding the role of predators in indirectly maintaining the integrity of the seaweed bed ecosystem.

The following questions were addressed in this study: 1) How does season affect urchin feeding rate and response to predation risk? 2) How does *H. crassispina* hunger state modify its foraging behavior in the presence of odor cues perceived to be a threat and a non-threat? and 3) How does urchin hunger state affect their predator avoidance behavior in the field?

## 2. MATERIALS AND METHODS

### 2.1. Collection and maintenance of urchins and algae

Purple urchins (*Heliocidaris crassispina* Agassiz, 1864) were collected from the coastal waters of Kashiyama Town, Nagasaki Prefecture, Japan. Urchins were brought to the Institute for East China Sea Research, Nagasaki University, approximately 3.7 km south of the collection site. Urchins were placed inside an outdoor one-ton tank (170 × 110 × 70 cm) with a constant flow of sand-filtered seawater and aeration. A Tidbit v2 (Onset Corp.) temperature logger monitored the ambient water temperature. Urchins were fed *ad libitum* with an assortment of fresh algae collected from Omura Bay, Nagasaki, Japan. The feeding experiments used *Sargassum patens* C. Agardh and were also collected from Omura Bay. Stock *S. patens* were kept in a separate outdoor tank which received water overflowing from the urchin stock tank. Urchins were acclimatized to ambient laboratory conditions for one week prior to the experiments (ambient temperature range for summer: 22.7 ± 4.79 °C and winter: 15.9 ± 3.52 °C; mean ± SD). Experiments involving Summer-Autumn and Winter-Spring seasons are hereafter referred to as Su-Au and Wi-Sp, respectively.

Urchins were starved by placing selected individuals in a separate container with no food for one week prior to experiments. This allowed for standardization of their nutritional condition and to elicit a stronger hunger response prior to experiments (Scheibling & Anthony 2001).

### 2.2. Laboratory experiment 1: Urchin grazing rate

To test the hypothesis that the ambient *H. crassispina* grazing rate was influenced by temperature and season, a flow-through rectangular tank (70 × 112 × 12 cm) was prepared. Ten numbered containers (2.96 L) were set in the tank, separated into two treatments. A feeding treatment that included urchins and algae and a control which contained only algae. A continuous water supply (11 L min^−1^) was provided by an overhead perforated polyvinyl chloride (PVC) frame. A Tidbit v2 (Onset Corp.) temperature logger recorded ambient water temperature.

A total of 24 trials (24 hours each trial) were conducted for both Su-Au (July-November 2018) and Wi-Sp (February-April 2019). Urchins used in Su-Au and Wi-Sp had test sizes of 4.28 ± 0.30 cm and 4.33 ± 0.45 cm (mean ± SD), respectively. A total of 144 urchins were used in both seasons. There were 4 control treatments and 6 feeding treatments for each trial. The purpose of the controls was to measure biogenic changes to the algae other than the effect of grazing. Whole *S. patens* thalli were removed of epiphytes and other debris and cut into portions. The cut portions were dried with paper towels and weighed to the nearest 0.1 g to obtain initial fresh weight. Urchins were weighed to the nearest 0.1 g while their horizontal test width was measured using a firm-joint outside-caliper and a Vernier caliper to the nearest 0.01 cm. *S. patens* cuttings and urchins were haphazardly assigned to containers. A mesh-net frame was placed over the tank to cover all containers to prevent urchins from escaping the container. At the end of each trial, the remaining uneaten algae were collected, dried with paper towels and re-weighed to obtain final fresh-weight. Urchin feeding rate was the difference between the final and initial weight with units g algae day^−1^.

### 2.3. Laboratory experiment 2: Effect of positive and negative chemosensory cues on urchin foraging behavior

The experiment was designed to test the hypothesis that urchins will modify foraging behavior when exposed to cues coming from dead conspecifics compared to controls (no odor cues).

The experiment used a flow-through tank similar to that of the previous experiment. However, water was supplied at a steady rate of 2.5 L min^−1^ through a hose. The hose was placed so that water flowed through the floor, in the middle of the tank. Water exited the tank through a 6 cm diameter hole in the tank wall, located 3 cm above the tank floor. Five concentric rings, 5 cm apart were marked on the tank floor around the water supply. The outermost ring defined the edges of the region of interest (ROI) where urchin behavior was recorded with a time-lapse camera (GoPro, Hero 4). At the center of the ring and above the hose, a perforated PVC cap was placed.

The experiment was started by placing one live urchin 10 cm from the center of the ROI. The camera was mounted 40 cm above the ROI and the field of view (FOV) included the entire tank (Supp. Figure 1). The images were recorded every 30 sec and experiment was conducted for 1 hr. Four treatments were defined, 1) a control (no dead urchin or algae; no chemosensory cue) 2) an algae treatment (algae only; positive chemosensory cue), 3) a dead urchin treatment (dead urchin only; negative chemosensory cue), and 4) a dead urchin and algae interaction treatment (combined chemosensory cues). To expose the test urchin to the treatment effect, algae were attached to the top of the PVC cap with clips while a recently crushed *H. crassispina* was placed within a mesh bag below the cap. Therefore, water flowing through the hose and through the cap ensured that chemosensory cues from the treatment would be dispersed outwards across the tank. The experiment was conducted on both urchin hunger states (i.e., starved and fed) and in Su-Au and Wi-Sp.

An opaque plastic sheet covered the entire experimental apparatus to remove all ambient light. However, below the sheet, a red LED lamp (ISL-150X150, CCS Inc.) provided enough light to record images while minimizing light disturbance to the urchins (Flukes et al. 2012). After every trial, the test urchins were removed, and the experiment chamber was rinsed thoroughly with freshwater and seawater to eliminate chemical cues from the previous experiment.

Experiments were conducted in the Su-Au (August-November 2018) and Wi-Sp (April-May 2019). A total of 111 individuals with test diameter 4.31±0.32 cm (mean ± SD) were used for the Su-Au experiment and 88 individuals with test diameter 4.27±0.58 (mean ± SD) were used for the Wi-Sp experiment.

For each trial, the time-lapse images were concatenated into an mpeg-4 video using FFmpeg (FFmpeg Developers 2018) at a frame rate of 10 frames per second (fps). Videos were analyzed with Tracker (ver.5.0.6; Brown 2018) to determine the movement pattern of the urchins. Each video frame (i.e., image) was counted as one event of a particular behavior. The following behaviors were possible: 1) None: Any immobile behavior within the region of interest. The urchin does not move up to 3 cm from its starting point. 2) Movement: The urchin is freely moving inside the region of interest. 3) Interaction: The urchin makes contact with the center of the region of interest which may or may not contain seaweed or dead urchin. The change in seaweed weight was not measured. 4) Outside: When the urchin went outside the region of interest. The x and y coordinates of the sea urchin was analyzed to determine the time spent by an urchin performing a particular behavior (minutes) as well as sea urchin movement speed (cm min^−1^).

### 2.4. Cue dispersal rate

The chemical cue plume was visualized and quantified using a 2% Fluorescein tracer-seawater solution as a proxy. A 3 mm diameter hose was attached to the tank floor so that the tracer was injected below the PVC cap and perpendicular to the water flow. The 50 ml of tracer was injected at a rate of 1.6 ml sec^−1^. Dispersal of the tracer was recorded on video for 1 hour. Three trials were conducted per treatment including control. The time for the tracer to reach the 10 and 20 cm ring was recorded and analyzed to determine if there were any differences among treatments.

### 2.5. Light measurement

The spatial homogeneity of the red light provided by the LED lamp was also assessed. Light was measured using five light loggers (HOBO MX2202 Temp/Light, Onset Corp.) that were placed on each ring to form a line. After the initial measurement, the line was rotated 30 degrees, for a total of four times. At every rotation, light was measured for one hour.

### 2.6. Field experiment: Effect of positive and negative chemosensory cues on urchin foraging behavior in the field

To examine the effects of chemosensory cues by food and dead conspecifics on sea urchin behavior under natural conditions, we prepared a site that was located at a depth of 4-5 m in a barren rocky area adjacent to natural stands of *Sargassum macrocarpum* in Arikawa Bay (32.988014 °N, 129.118638 °E), Nakadorijima Island, Nagasaki, Japan. A 2 m^2^ plot of flat rocky substrate was selected. For each experimental trial, a 2 m tall slotted angle-bar tripod frame with an approximately 1 m^2^ plan area was deployed. A time-lapse camera (TLC200 PRO, Brinno, Taiwan) enclosed in a custom acrylic housing was secured to the top of the frame. Images were taken every 30 sec for a total of 3 hours and stored as a video with a frame rate of 15 fps. The experiment was conducted during slack tide, when the tidal current was negligible. A velocity logger (Compact-EM, Alec Electronics Co.) and a water level logger (HOBO U20-001, Onset Corp.) were deployed 1 m from the experimental plot to record hydrodynamic conditions during the experiment.

A weighted plastic cage was placed in the middle of the 1 m^2^ experimental plot. Drift algae (i.e. *Sargassum spp., Dictyopteris* spp.) common during the experiment period was collected and clipped outside of the plastic cage. For each treatment, five trials were conducted. For the control, a single urchin was placed in direct contact with the algae until they attached. For the dead urchin and algae treatment, a recently killed *H. crassispina* was added inside the cage with the algae to determine whether urchins would be repelled. Urchins were killed just before the experiment started, by crushing their test. The experiments were conducted first on the fed and then on the starved urchins. A total of 20 individuals with test sizes of 4.72±0.68 cm (mean ± SD) were used for the field experiment.

Video was analyzed with Tracker (ver. 5.0.6) (Brown 2018) to track the urchins, however the tripod attracted small fish, which occluded the field of view. Additionally, during a number of days, the area experienced relatively high waves, which vibrated the tripod that resulted in poor quality images. Thus, only the initial and final position (stay or flee) of the urchin was noted after the 3 hr experiment period.

### 2.7. Data analyses

#### Laboratory experiment 1: Urchin grazing rate

The urchin grazing rate data was analyzed using a Bayesian generalized linear model where the mean grazing rate (g algae urchin^−1^ day^−1^) was the response variable and the explanatory variable was the season. Weakly informative priors were used for the intercept and coefficients. The gaussian distribution with a location of 1.1 and a scale of 2.5 was the prior for the intercept, a gaussian distribution with a location of 0 and scale of 2.5 was the prior for the coefficients, and an exponential prior with a rate of 1 was the prior for the error term.

#### Laboratory experiment 2: Effect of positive and negative chemosensory cues on urchin foraging behavior

Initial inspection of the urchin movement behavior data revealed an over-abundance of zeroes, as not all behaviors were represented equally for every trial. Meanwhile, some behaviors had more occurrences compared to others. Both observations generally cause issues such as zero-inflation and over-dispersion. To overcome this, hurdle-models were used for model fitting because of its two-step procedure beginning with a Bernoulli probability, which evaluates whether a count is non-zero. Next, if a positive, non-zero value was found, this “hurdle” is crossed, and then proceeds with a truncated-at-zero count distribution model for the non-zero state (Lewin et al. 2010, Kassahun et al. 2014). This is similar to a decision-making process because the outcome of an individual’s behavior can depend on existing environmental conditions (i.e. treatment).

Specifically, the urchin behavior in the laboratory experiments were analyzed with a hurdle-negative binomial model (Eq. 1),

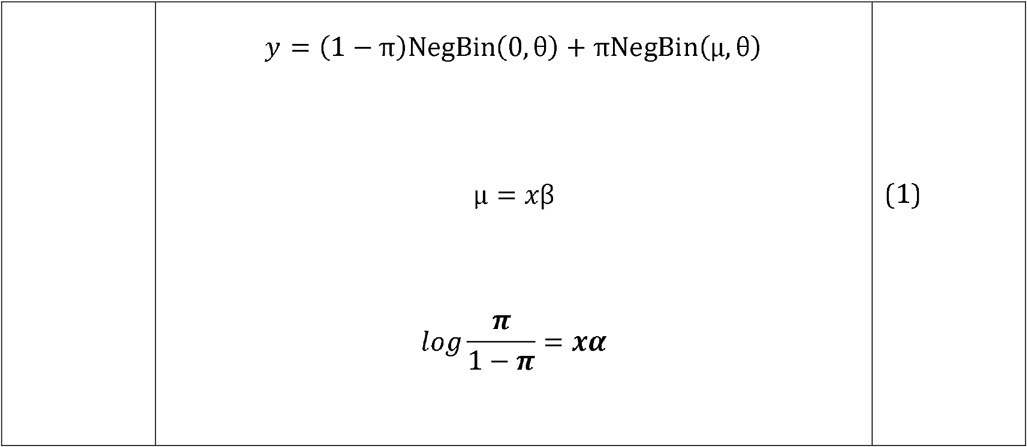

where, *y* is a vector of observations, and in this case the number of occurrences for a behavior during the 1-hour observation period. π is a vector of probabilities for non-zero values, α and β are vectors of coefficients for a model including all treatment interactions. *x* is a matrix of factors that include all treatment interactions. The number of occurrences is assumed to follow a negative binomial distribution, with a vector of locations μ and a scale θ. The main treatments are the presence or absence of algae and the dead urchin, the hunger state of the test urchin, season (i.e., Su-Au or Wi-Sp), and the type of behavior, excluding the behavior “none” (see Fletcher et al. 2005, Zuur et al, 2009).

In the case of the sea urchin speed and time spent per behavior, where the response was a continuous variable, a hurdle-gamma model was applied. The structure of the model is similar to Eq. 1, however rather than a negative binomial distribution a gamma distribution is assumed. In this case *y* = (1-π)Γ(0, θ) + πΓ(μ, θ). For more details on the merits of the hurdle model, see Lewin et al. 2010.

The β coefficients of the all hurdle models were given weakly informative Student’s t-distributions as prior distributions, with 3 degrees of freedom, a location of 0, and a scale of 1. The α coefficients were given logistic distributions as priors with a location of 0 and a scale of 1. The prior for θ was a Γ distribution with a shape and scale of 0.01.

#### Cue dispersal rate

The data on the Fluorescein tracer dispersal experiment was analyzed using a Bayesian generalized linear model where the time it took for the tracer to reach the 10 cm mark was the response variable while the explanatory variables were the experimental treatments. The prior distributions were similar to that of the urchin grazing rate analysis, however the location for the prior intercept was 18.

#### Light measurement

For the light experiment, a Bayesian generalized linear model was fitted to the data where the response variable was the light level and the explanatory variables were the position of the light loggers. The prior distributions were similar to that of the urchin grazing rate analysis, however the location for the prior intercept was 1.7.

#### Field experiment: Effect of positive and negative chemosensory cues on urchin foraging behavior in the field

The field experiments were analyzed with a Bayesian binomial generalized linear model with a random intercept for the tidal state (Eq. 2).

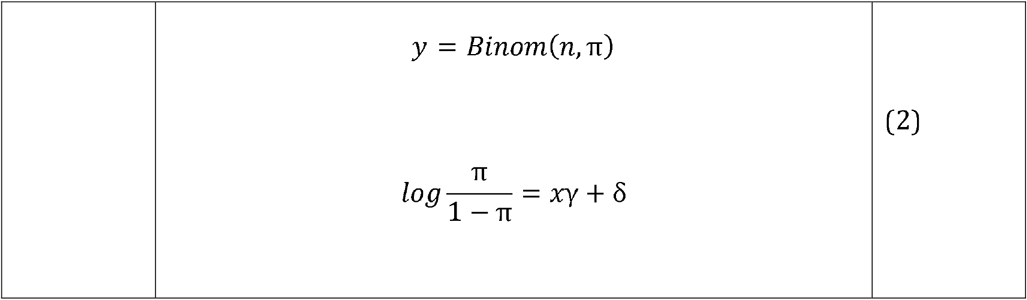

In Eq. 2, the *y* is the vector of observations, *n* is the vector of total trials, and π is the vector of probabilities. *x* is the matrix of treatments, which in this case is a linear combination of the hunger state of the test urchin (i.e., starved or fed) and the presence or absence of the dead urchin. δ is a random intercept for tidal state. The vector of coefficients is γ. The prior distributions for the coefficients and random intercept was a Student’s t-distribution with 3 degrees of freedom, a location of 0, and a scale of 5.

All statistical analyses were done using R version 3.6.1 (R Development Core Team 2019) and all Bayesian inference was done with Stan (Stan Development Core Team 2019) through the brms (Bürkner 2017) and RStanarm packages (Goodrich et al. 2018). Stan primarily uses a Hamiltonian Monte Carlo Sampler to construct the posterior distributions of the parameters. For all models, a total of four chains were evaluated to generate 2000 samples per chain. All chains of all models were assessed for convergence.

## 3. RESULTS

### 3.1. Laboratory experiment 1: Urchin grazing rate

The results revealed differences in feeding rates between seasons. Sea urchins had higher expected mean feeding rates in Su-Au of about 1.3 g algae urchin^−1^ day^−1^ (1.2-1.5 95% HDI) (Table 1: A). Conversely, sea urchin feeding rates decreased to 0.8 g algae urchin^−1^ day^−1^ (0.6-1.0 95% HDI) during Wi-Sp. There was a 23% difference in the mean maximum ambient water temperature between seasons (Figure 1).

**Fig. 1.**
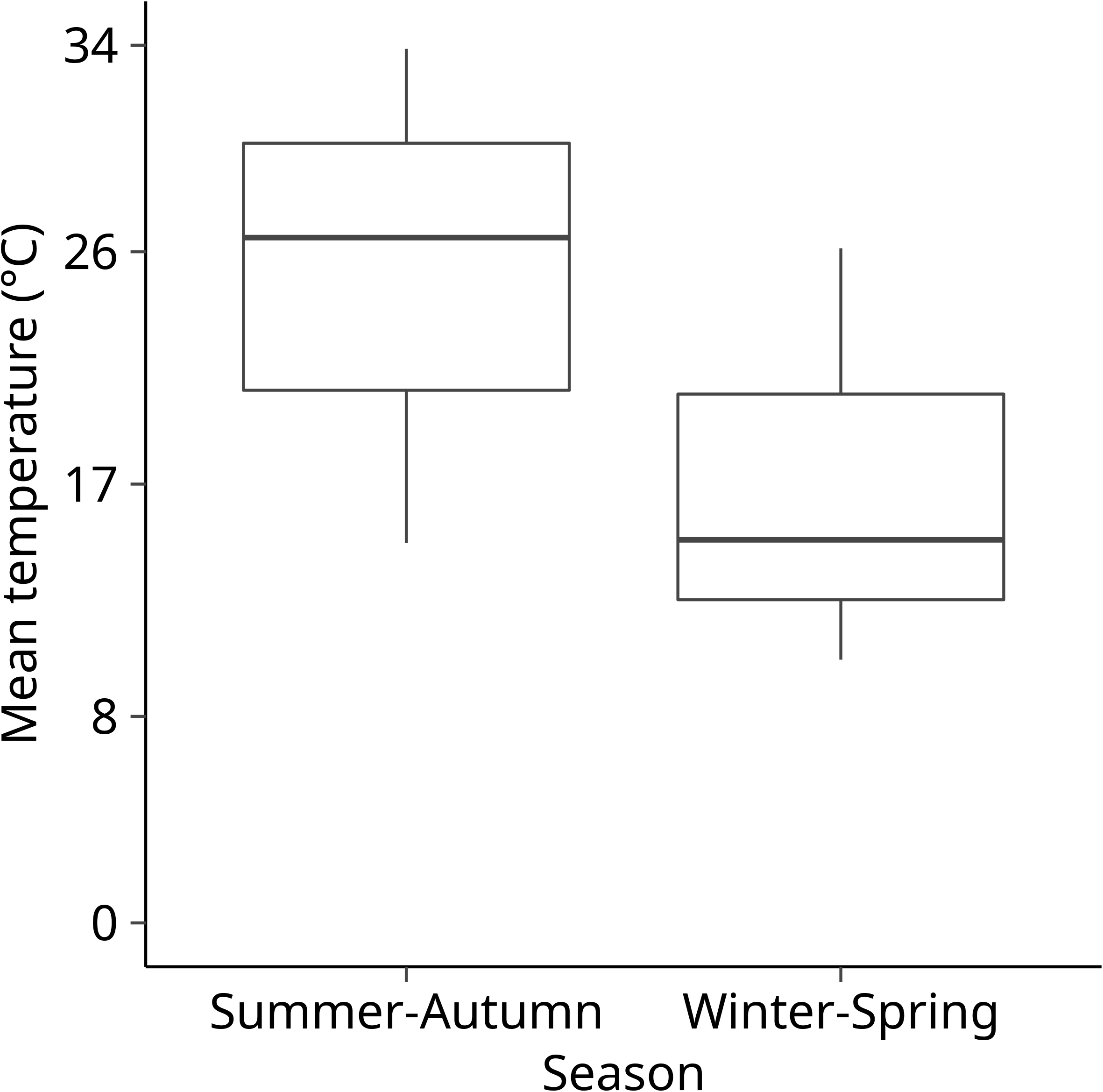
Mean temperature difference between Summer-Autumn and Winter-Spring seasons in the feeding rate experiment.

**Table 1.** Results of the Bayesian generalized linear models on A) Experiment 1: feeding rate and season, B) cue dispersal time to reach the 10 cm mark. The table shows the estimates, expected value and the lower and upper limits of the 95% highest density interval (HDI) of the expected value.

| Estimates | Expected value | 2.5% | 97.5% |
| --- | --- | --- | --- |
| **A.) Lab experiment 1: Feeding rate (g algae urchin<sup>-1</sup> day<sup>-1</sup>)** |  |  |  |
| Summer-Autumn | 1.3 | 1.2 | 1.5 |
| Winter-Spring | 0.8 | 0.6 | 1.0 |
| **B.) Cue dispersal time (sec)** |  |  |  |
| Control | 12.2 | -1.4 | 24.9 |
| Algae | 17.5 | 4.3 | 31.1 |
| Dead urchin | 23.0 | 9.8 | 36.5 |
| Algae + dead urchin | 19.7 | 7.8 | 33.4 |

### 3.2. Laboratory experiment 2: Effect of positive and negative chemosensory cues on urchin foraging behavior

#### 3.2.1. Behavior counts

Sea urchin activity was not discernibly affected by the light intensity (Supp. table 1) throughout the experiment. The time-lapse experiment showed that the counts of the four behavior types varied widely across sea urchin condition, season, and treatment. The occurrence of the immobile behavior “none” occurred more in Wi-Sp (starved: 15 urchins, fed: 21 urchins) than in Su-Au (starved: 16 urchins, fed: 15 urchins) (Figure 2). The occurrence of this behavior in Wi-Sp represents 40.9% of the sea urchins used in that season while this behavior represented about 27.9% of the total sea urchins used in the Su-Au.

**Fig. 2.**
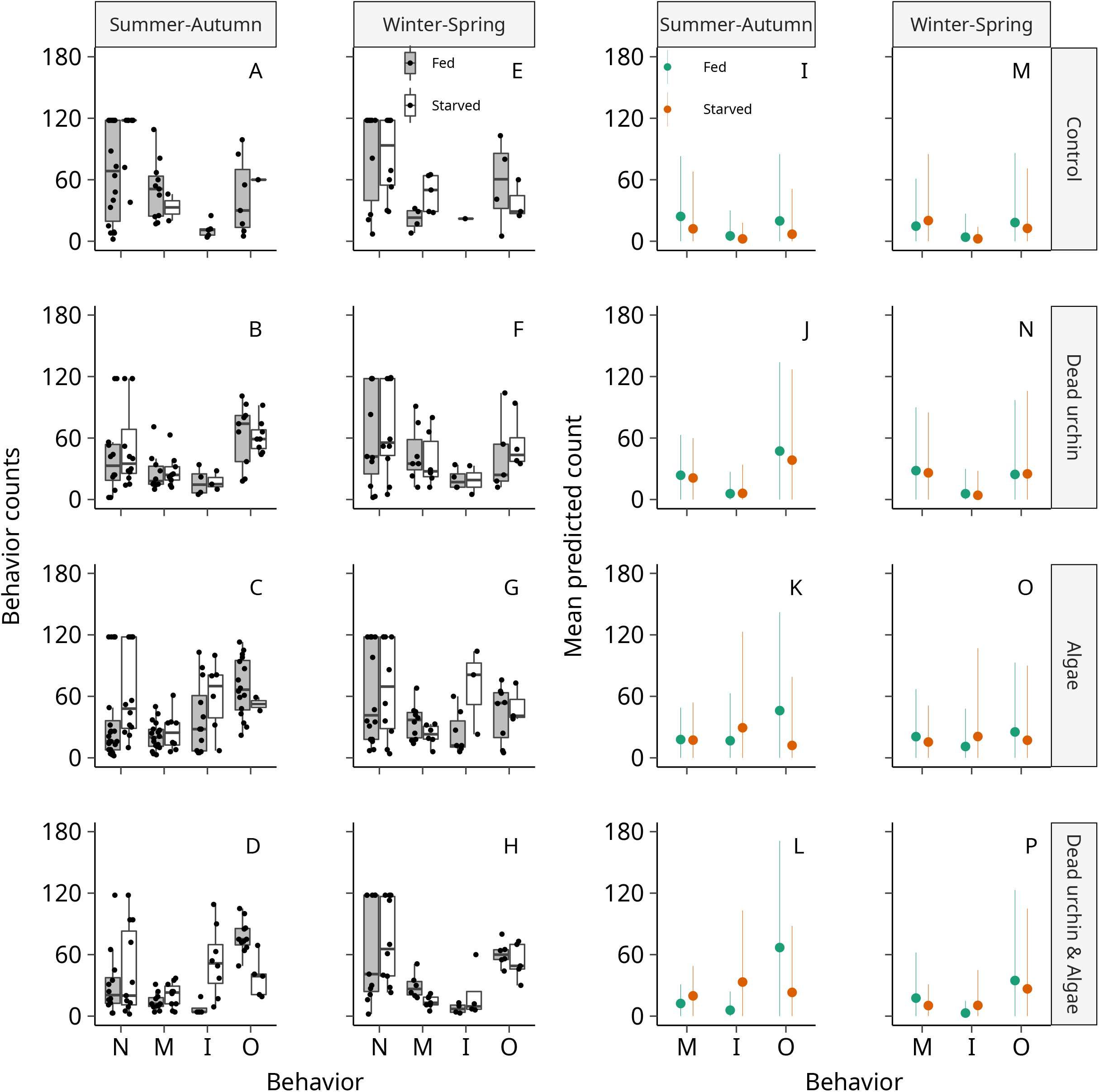
Experiment 2 empirical data (A to H) and model prediction (I to P) of the count of urchin behaviors. The columns indicate the seasons and the rows indicate the treatments. Behaviors are abbreviated as N (none), M (movement), I (Interaction), and O (Outside). The extents of the boxplots indicate the 25% and 75% percentiles and the horizontal line indicates the median. The whiskers extend 1.5 times the inter-quartile range and the overlaid points are the observations for each behavior. In subfigures I to P, the points indicate the predicted mean while bars are the 95% highest density intervals (HDI) of the predictions.

Sea urchins were generally more active in Su-Au than in Wi-Sp season. For interaction, this behavior was more frequent among trials in Su-Au (6%) than in Wi-Sp (2%). Interaction was also more frequent among starved sea urchins (5%) than for fed urchin (4%).

Comparing the effects of algae only and combined chemosensory cues treatment shows differing responses across nutritional states. The model shows that when only an algae is present, fed urchins had a mean interaction count of up to 33.8 (2-71 95% Highest density prediction interval, HDI) during Su-Au and 26.6 (1-61 95% HDI) in Wi-Sp. When a dead urchin was present together with the algae (Figure 2L, P), this led to a mean decrease in their interaction counts to 12.9 (1-32 95% HDI) and 8.3 (1-21 95% HDI) during Su-Au and Wi-Sp, representing about 61.8% and 68.7% decrease, respectively. For starved urchins, their hunger state led to high interaction counts relative to fed urchins during Su-Au, 66.7 (1-161 95% HDI) and 66.0 (1-163 95% HDI) for Wi-Sp when only algae was present. Under the combined chemosensory cues treatment, starved urchins had interaction counts of to 51.1 (5-116 95% HDI) during Su-Au and 24.5 (2-60 95% HDI) in Wi-Sp. This shows a 23.4% and 62.9% decrease between seasons, respectively. In Wi-Sp, starved urchins also had a higher proportion of immobile individuals across both hunger states. The presence of the dead urchin with the algae also increased the number of “outside” behaviors across both seasons for fed urchins (8.7% and 25.2% for Su-Au and Wi-Sp, respectively), but not for starved urchins. They show decreased “outside” behaviors of up to 33.2% and 6.11% for Su-Au and Wi-Sp, respectively. Overall, both hunger states seem to be sensitive to the chemical cues from dead urchins but starved urchins appear to interact more with the algae despite predation cues. The expected value and prediction intervals for behavior counts are shown in Suppl. Table 2 while the probability of behaviors achieving zero counts are shown in Suppl. Table 5.

#### 3.2.2. Time spent per behavior

The time-lapse experiment demonstrated the ability of the treatments, condition and the season to influence the sea urchin’s allocated time performing a specific behavior (Figure 3). Overall, 34% of sea urchins spent the entire 1-hr experiment period immobile. Of that number, 42% and 27.9% occurred during Wi-Sp and Su-Au experiments, respectively.

**Fig. 3.**
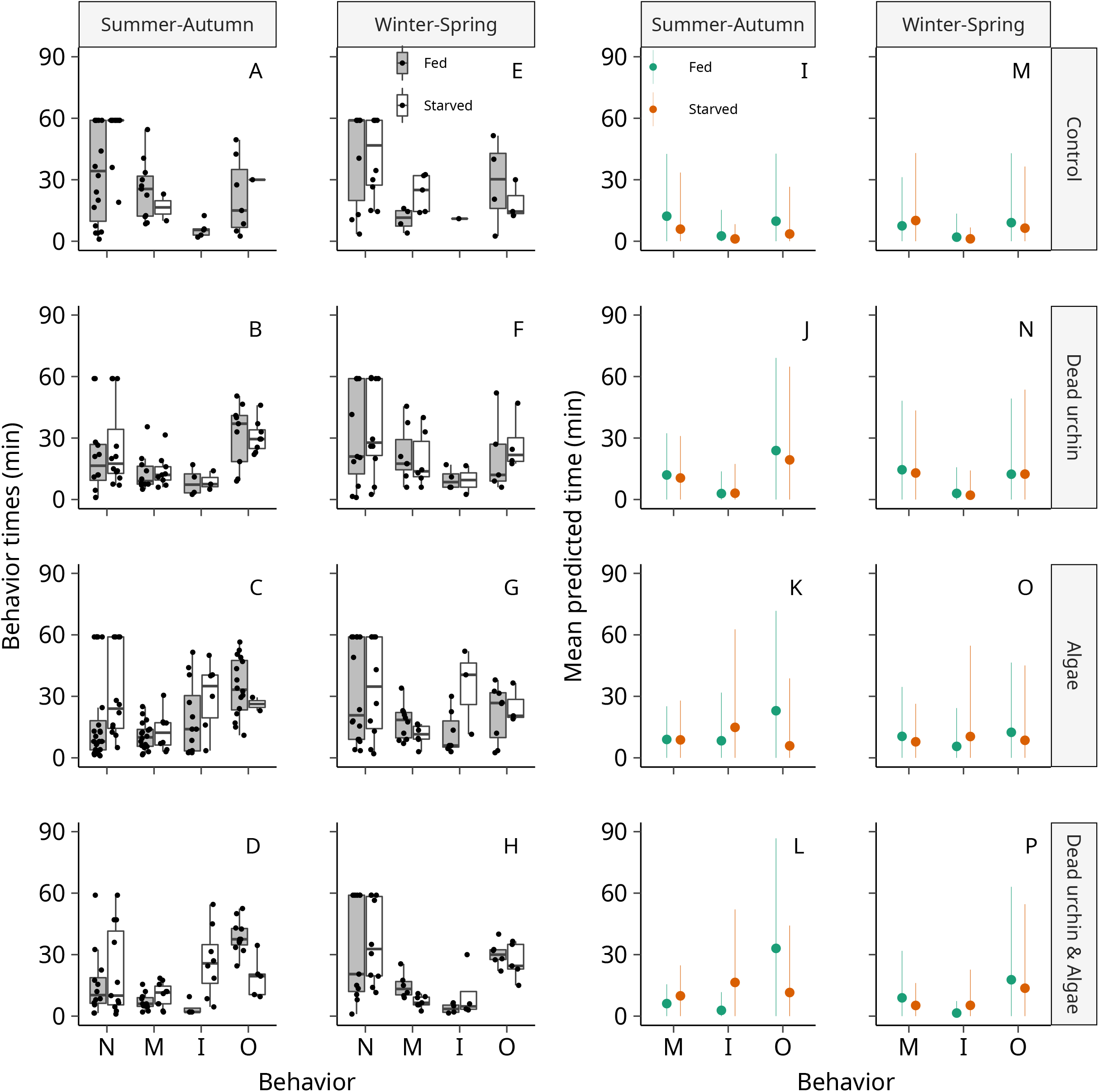
Experiment 2 empirical data (A to H) and model prediction (I to P) of the time urchins spent performing a behavior (min). The columns indicate the seasons and the rows indicate the treatments. Behaviors are abbreviated as N (none), M (movement), I (Interaction), and O (Outside). The extents of the boxplots indicate the 25% and 75% percentiles and the horizontal line indicates the median. The whiskers extend 1.5 times the inter-quartile range and the overlaid points are the observations for each behavior. In subfigures I to P, the points indicate the predicted mean while bars are the 95% highest density intervals (HDI) of the predictions.

The model predictions show that the presence of a dead urchin had an influence over the time spent urchins were performing a particular behavior. Under the algae treatment, fed urchins had an average interaction time of about 17.7 min (0.988-40.5 95% HDI) in Su-Au and 13.6 min (1.24-33.3 95% HDI) in Wi-Sp. Under the combined chemosensory cues treatment, fed urchins had a mean interaction time of 6.5 min (0.180-14.7 95% HDI) in Su-Au and 4.3 min (0.180-10.6 95% HDI) in Wi-Sp. This shows an 11.2 and 9.3 minute difference in interaction time across seasons, respectively. Starved urchins were predicted to have relatively higher mean interaction times relative to fed urchins when only algae was present (Su-Au: 33.4 min, 1.61-82.1 95% HDI; Wi-Sp: 33.2 min, 0.982-81.1 95% HDI). Under the combined chemosensory cues treatment, urchins in Su-Au had mean interaction time of 25.3 min (0.929-59.2 95% HDI) while urchins in Wi-Sp had a mean interaction time of 12.9 min (0.552-31.5 95% HDI). This shows a decrease of 8.1 and 20.3 min for Su-Au and Wi-Sp, respectively. The time spent outside the ROI also increased across both seasons for fed urchins (4.1 and 7.6 minutes for Su-Au and Wi-Sp, respectively). The starved urchins show decreased time outside the ROI by about 8.4 and 0.9 minutes for Su-Au and Wi-Sp, respectively. As expected, when a dead urchin was present, the fed urchins interacted less with the algae and increased their time spent outside, indicating that the urchins were repelled by the presence of the dead urchin chemical cues. Similarly, starved urchins show a decrease in interaction time but by a slightly lesser rate. Their decrease in time spent outside despite the presence of the dead urchin suggests that the hunger state was able to influence urchin behavior. Estimates and prediction intervals for time spent per behavior are shown in Suppl. Table 3 while the probability of the behavior time becoming zero minutes are shown in Suppl. Table 6.

#### 3.2.3. Urchin movement speed

The time-lapse experiment revealed urchin movement speeds varied across the treatments depending on their condition and season. It was noted that even when the urchins were exhibiting the behavior “none”, small movement speeds were recorded as urchins were shuffling in place to within the 3cm limit. The urchins exhibited greater speeds when a dead urchin was present. Overall, fed urchins had higher move speeds relative to the starved urchins (Figure 4).

**Fig. 4.**
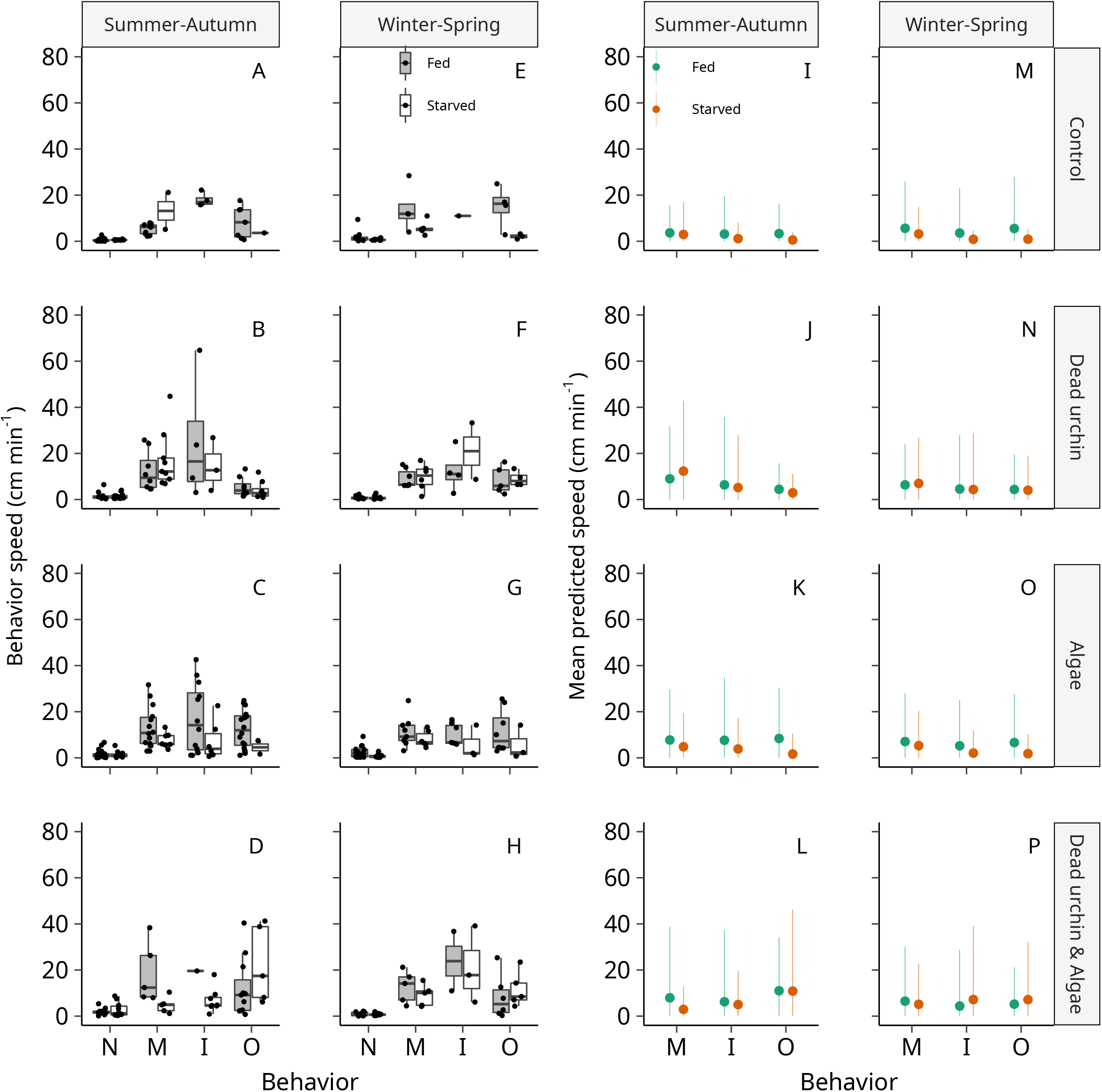
Experiment 2 empirical data (A ot H) and model prediction (I to P) on urchin speed (cm min^−1^) for each behavior. The columns indicate the seasons and the rows indicate the treatments. Behaviors are abbreviated as N (none), M (movement), I (Interaction), and O (Outside). The extents of the boxplots indicate the 25% and 75% percentiles and the horizontal line indicates the median. The whiskers extend 1.5 times the inter-quartile range and the overlaid points are the observations for each behavior. In subfigures I to P, the points indicate the mean while bars are the 95% highest density intervals (HDI) of the predictions.

The model predictions indicate that urchins tend to move at a different pace depending on the treatment (Figure 4). When only the algae was present, fed urchins had mean interaction speeds of 16.6 cm min^−1^ (0.243-46.5 95% HDI) in Su-Au and 12.5 cm min^−1^ (0.164-34.3 95% HDI) in Wi-Sp. Under the combined chemosensory cues treatment, fed urchins had mean interaction speeds of 24 cm min^−1^ (0.141-65.9 95% HDI) in Su-Au and 21.1 cm min^−1^ (0.375-64.4 95% HDI) in Wi-Sp. Between treatments, fed urchin speed while interacting with the algae increased by 7.4 and 8.6 cm min^−1^ during Su-Au and Wi-Sp, respectively, when a dead urchin was present. For starved urchins, under the algae only treatment, predicted interaction speed was 7.9 cm min^−1^ (0.030-21.7 95% HDI) in Su-Au and 6.8 cm min^−1^ (0.073-18.7 95% HDI) in Wi-Sp. In the combined chemosensory cues treatment, interaction speeds were 8.3 cm min^−1^ (0.043-23.0 95% HDI) in Su-Au and 21.4 cm min^−1^ (0.302-60.9 95% HDI) in Wi-Sp. There seems to be a slight increase in urchin speed during Su-Au by about 0.4 cm min^−1^, but a large rate of increase of about 14.6 cm min^−1^ for urchins in Wi-Sp. Movement rates within the ROI also increased for fed urchins while outside speeds increased for starved urchins. Examining urchin speeds show that signals of predation may cause stress to *H. crassispina* as indicated by the relatively high movement speeds across both nutritional states and seasons even when outside the ROI. Estimates and prediction intervals for urchin speeds per behavior are shown in Suppl. Table 4 while the probability of behaviors becoming zero cm min^−1^ are shown in Suppl. Table 7.

#### 3.2.4. Cue dispersal rate

The results from the experiment on the rate of spread of the Fluorescein tracer dye showed high variation among the trials (Table 1: B). The time it took for the tracer to reach the 10 cm mark was modelled since it represented the area where the urchin would first encounter the chemosensory cues coming from the center of the ROI. The control, with nothing beneath and above the treatment container, took the least amount of time and had an expected mean time of 12.2 sec (−1.4-24.9 95% CI). The treatments could be ranked from those that took the least amount of time to the greatest amount of time and resulted in an order of control, algae effect, dead urchin and algae interaction effect, and dead urchin effect. The wide range for all treatments was due to a low sample size (3 trials per treatment). However, high variations between trials among the algae, dead urchin and dead urchin and algae treatments suggested that the variation was associated with the size of the dead urchin or the density of the algae used as treatment for the experiment.

### 3.3 Field experiment and environmental conditions: Effect of positive and negative chemosensory cues on urchin foraging behavior in the field

The field experiment showed that sea urchin condition produced discrete responses between starved and fed urchins to the presence of dead conspecifics adjacent to an available food source. Of the 20 sea urchins used in the experiment, all 10 starved sea urchins (100% of the starved condition) stayed and remained in contact with the treatment cage. For the fed sea urchins, only 3 stayed (30%) while 7 fled from the treatment cage (70%). Of the 7 sea urchins which fled, 4 sea urchins (40%) were from the treatment which contained the dead urchin. The binomial model predictions suggested a strong link between urchin condition and outcome of behavior (Figure 5).

**Fig. 5.**
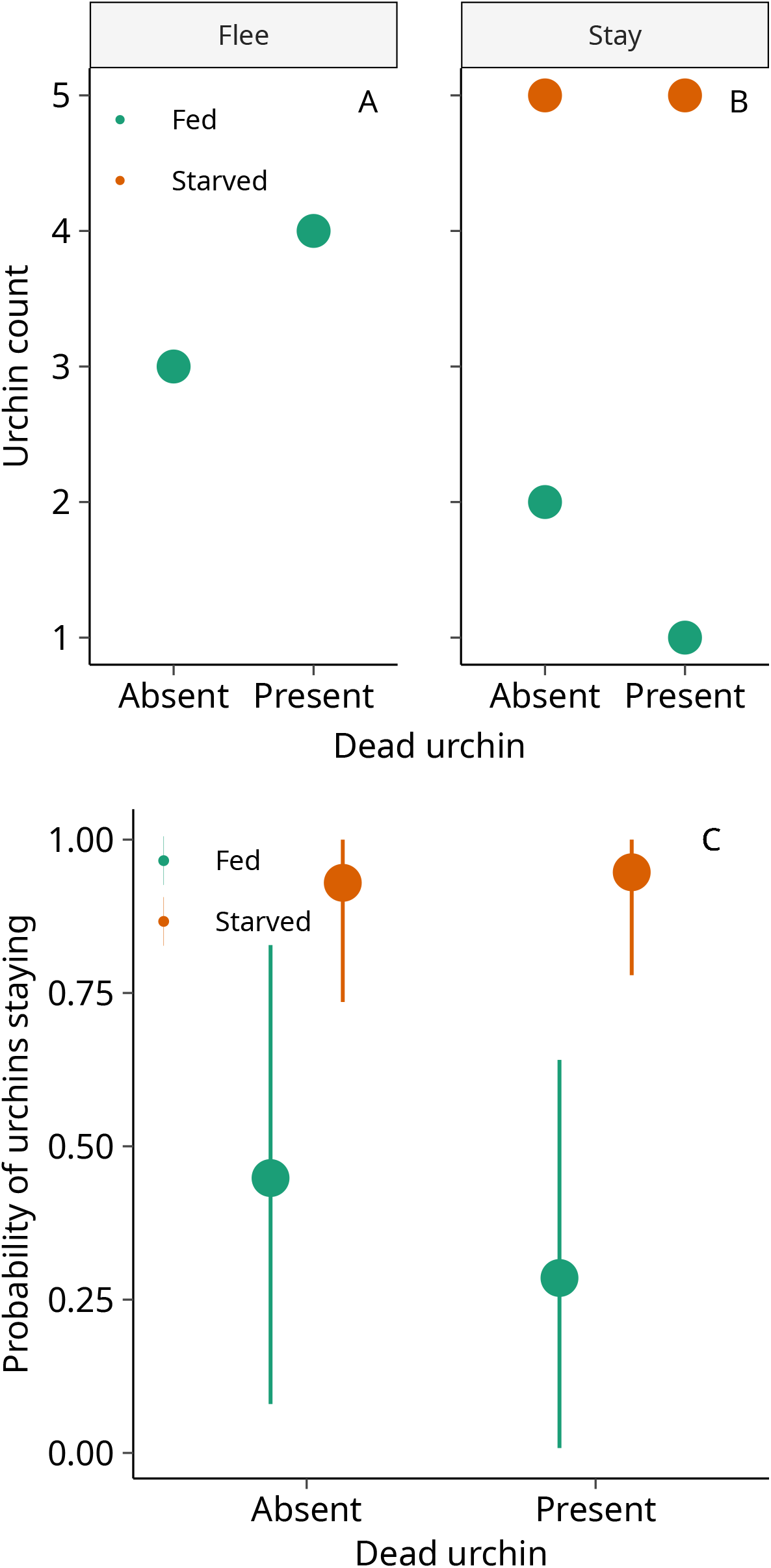
Field experiment empirical data (A, B) and binomial model prediction (C) on urchin behavior outcome in the field. For A and B, the y-axis shows the number of urchin counts, the x-axis shows either the presence or absence of a dead urchin with the algae as treatment and the columns show the response of the urchins. For the results of the binomial model, the y-axis shows the binomial probability of urchin behavior (i.e. flee: 0, stay:1). The x-axis shows the presence or absence of a dead urchin together with the algae. Points show the expected means while bars are the 95% highest density intervals (HDI) of the expected values. Results show that starvation makes urchins less likely to flee regardless of the presence or absence of a dead urchin.

Of the 20 trials conducted, 11 experiments were conducted during low slack tide while 9 were conducted during high slack tide. In general, mean temperatures and mean current speeds were higher during low tide relative to high tides (Supp. Figure 2).

## 4. DISCUSSION

### 4.1. Factors affecting sea urchin behavior patterns

The result of our study provides evidence that trait-mediated indirect interactions (TMII) is an effective component of top-down trophic cascades (Schmitz et al., 2004). We found discrete behavior patterns between starved and fed *H. crassispina*, suggesting that hunger-state determines an individual’s propensity to accept a certain degree of risk to acquire critically needed resources. For starved urchins, there was a greater proportion of urchin interaction behavior and increased interaction time with the algae despite the presence of a dead urchin. Fed urchins exhibited predator avoidance behaviors, observed as decreased interaction and increased occurrences of behaviors spent outside the ROI when a dead urchin was placed together with the algae.

Studies on predator-prey relationships highlight the “Hobson’s choice” (i.e., face the risk of predation or starve) dilemma all prey species face upon venturing out from the safety of their refuge when they forage for food (Clark 1994). Ultimately, the decisions prey species make lean towards optimizing the trade-off to their advantage by minimizing risk while maximizing benefits. However, intrinsic (i.e., reproductive condition, and hunger level) and extrinsic (i.e., temperature, light, and salinity) factors also play an important role in affecting decision-making processes for aquatic organisms. For example, the effects of starvation in urchins not only impacted their energy reserves but also their gut and gonad indices (Lawrence 1970). A study on the effects of starvation on *H. crassispina* and *Hemicentrotus pulcherrimus* showed that gut clearance was achieved in 3 days for *H. crassispina* and 6 days for H. pulcherrimus. Additionally, H. pulcherrimus survived a maximum of 49 days without food albeit negatively impacting gut and gonad indices (Kaneko et al. 1981 in Agatsuma et al 2013). In our study, the one-week starvation period induced a hunger response which appeared to override predator avoidance behaviors. The starved urchins, which had an energy deficit, were willing to accept greater risks by feeding longer and more frequently in the presence of a dead urchin to increase energy reserves, hence supporting the asset protection principle (Clark 1994). It should be noted that the microcosm experiment utilized a relatively small chamber which may have allowed a faster saturation of sea urchin effluents and thus increase the urchin responses artificially. Our experiment did not provide refuge for urchins which may explain their rapid movements inside and outside the ROI when dead urchin cues were present. Furthermore, the manipulation of sea urchin condition by starvation as done in our study does not fully mimic conditions in the field. Recall that sea urchins are generalist algal feeders (Vadas 1977) and have flexible dietary preferences and are omnivorous (Rodriguez-Barreras et al. 2015). *H. crassispina* in seaweed bed habitats generally had higher gonad indices and were larger in size compared to urchins collected from a habitat dominated by *Corallina* spp. (Yatsuya & Nakahara 2004a). In the barrens, they were more cryptic and switched to feeding on a mixed diet composed of drift Sargassum and calcareous algae (Yatsuya & Nakahara 2004b). Other signs of food limitation manifested themselves as differences in morphometric changes such as smaller than average test size and longer jaw lengths to maximize grazing efficiency (Pedersen & Johnson 2008).

A recent study utilizing starved and satiated *Strongylocentrotus droebachiensis* showed that both sea urchin groups did not react adversely to the presence of a live nearby predator (*Cancer borealis*) (Harding & Scheibling 2015). Their findings indicate that the olfactory cues coming from live predatory crabs did not reduce urchin foraging behavior in the laboratory and in the field. In contrast, studies on chemical alarm cues showed dead conspecifics and chemically-labelled predators had a strong adverse effect on urchin behavior (Campbell et al. 2001, Morishita & Barreto 2011). Specifically, the urchins distinctly avoided waters conditioned with gut, coelomic, and gonad homogenates, which were the materials most likely to be exposed when a predator breaks an urchin’s test (Campbell et al. 2001).

In our study, instead of completely avoiding the source of the chemical cues coming from the dead urchin treatment, some urchins actually approached the dead urchin treatment. In small prey species of fish, this behavior is known as predator “inspection”, and had distinct importance for prey species because this functions as a learning tool to enable naive prey to associate predators with danger (Magurran & Girling 1986). As prey grow and reach sizes which act as refuge from direct predation, their fear of predators remain and continue to affirm the effects of top-down control (Pessarrodona et al. 2019). For urchins, since olfaction occurs when odor molecules reach receptors in their tube-feet, predator inspection may need to occur at close range as odor molecules increase in concentration. The next time they encounter familiar chemical cues relating to risk of predation, they may better assess the motivation of the predator and the relative risk of an impending predation event (Clark 1994). Furthermore, for urchins living in urchin barrens, it may be possible that these urchins had reached large population sizes due to the absence of their natural predators. The absence of predators meant that it was likely that they were naive and had little chance in encountering chemical cues relating to predation.

When comparing sea urchin behavior patterns across seasons, we found that there was a discrete pattern observed between the Su-Au and Wi-Sp experiments. In general, sea urchins were more active and exhibited higher speeds during Su-Au compared to sea urchins used in Wi-Sp. The greater decrease in the interaction frequency and interaction time in Wi-Sp for starved urchins was attributed to lesser urchins interacting with the treatment as well as more urchins moving outside. Interestingly, urchin speed was predicted to be highest in Wi-Sp when starved individuals were exposed to dead urchins together with food. This was likely to be an evasive behavior in response to the scent of the dead urchin since the proportion of outside behaviors and movement speeds also increased. At the same time, when only food was present, starved urchins interacted with the algae longer and moved slower, indicating a stronger intent to feed, compared to fed urchins (Figure 3). Their level of activity was also reflected in their feeding rates as urchins in August had the highest average feeding rates while urchins in February had the lowest rates (Table 1:A). This is a similar pattern found among cold-water urchin species where temperature was one of the main drivers of metabolic activity (Agatsuma et al. 2000, Brockington & Clarke 2001).

Studies on the reproductive biology of *H. crassispina* showed that this species had a distinct seasonal cycle in terms of gonadal development and maturation. In Nagasaki, Japan, a study on the reproductive patterns of *H. crassispina* (Yamasaki & Kiyomoto 1993 in Agatsuma et al 2013) has found that this species spawns during the months of July to August while their recovery period was from September to January. The rest of the following months were dedicated to growth and maturation of the gonads. This pattern was also similar with studies elsewhere in Japan (Kyoto: Yatsuya & Nakahara 2004a, Oga Peninsula: Feng et al. 2019) and in Korea (Yoo et al. 1982). In Hong Kong, a 7 to 8 month spawning period was recorded. This relatively long spawning period was represented by two distinct spawning events in May-June and September-October (Urriago et al. 2016). After every spawning event, urchins experienced an abrupt decrease in gonad indices as well as lipid and fatty-acid profiles (Martinez-Pita et al. 2010, Diaz de Vivar et al. 2019). The lipid and nutrient deficient state indicated that the urchins were in a low nutritional condition (Lawrence 1970). Urchins compensated by increasing their feeding rates beginning from the end of summer until next spring, coinciding with winter macroalgal blooms (Kaehler & Kennish 1996). Increasing feeding rates from summer ensured the accumulation of energy to support gonadal growth and maturation as reflected from the biochemical composition and other intrinsic gonad properties (Rocha et al. 2019). Hence, the rise in summer metabolic activity in urchins was only partially explained by temperature but was likely predominantly driven by feeding, growth and reproduction (Brockington & Clarke 2001).

### 4.2. Field experiment

The results provide evidence of urchin condition affecting the strength of behavior modification in the field. Compared to fed sea urchins, all starved urchins stayed regardless of the presence or absence of a dead urchin. The result of our experiment was in contrast with the field experiment using live crab predators where only a 6% flee response rate was recorded (Harding & Scheibling 2015). Few studies have previously investigated effects of predation risk cues on prey species in the field because of the inherent difficulty of controlling for local flow conditions. The data recorded from the field shows high variability in flow speeds as well as temperature between low and high tides. A laboratory study simulating the flow of chemical odor plumes in turbulent conditions suggests that the success of odor-guided navigation was greatly dependent on dilution and degree of shear-induced mixing of odor signals (Webster & Weissberg 2001). This is particularly true for small benthic invertebrates because sampling the water for odor molecules occurs at a relatively faster rate (Zimmer & Butman 2000) but at a lower height relative to the substrate (Smee & Weissberg 2006). Furthermore, organisms attempting to orient themselves relative to the direction of the odor-plume would find it challenging because odor dispersal occurs as intermittent odor packets interspersed with clean water (Finelli et al. 1999). In the present study, the sea urchins would have had no problem detecting the odor from the dead urchin and seaweed because they were placed in direct contact with the treatment cage at the start of the experiment, unlike in the laboratory experiment. Although concentrations of urchin effluent were not tested, it may be plausible that the immediate area surrounding the treatment cage would have been saturated with the dead urchin effluent. The fleeing response of some of the fed sea urchins appear to be a behavior related to minimizing predation risk in lieu of feeding opportunity. However, with only a short 3-hr experiment period, we were not able to observe how the starved urchins would behave once they have adequately fed on the seaweed or how long the dead urchin effluents remained effective. Hence, for future studies, we propose a longer observation time for urchin behaviors and identification of the components responsible for urchin alarm response and their maximum length of efficacy (i.e. Spyksma et al. 2020) as affected by dilution.

## 5. CONCLUSIONS

The behavioral responses of *Heliocidaris* crassispina in our experiments appear to encompass all classifications in the prey behavior types proposed by Fraser & Huntingford (1986). Our results showed that *H. crassispina* foraging behavior was flexible and was able to assess and adjust accordingly to the presence of chemical cues associated with predation. To some extent, our results support the idea that the type of season and phenology appear to modulate urchin behavior and foraging activity (Luttberg et al. 2003). Our experiments also demonstrate that the presence of a dead urchin does not prevent live urchins from interacting with the seaweed but instead decreases the interaction frequency, length of interaction time and increases movement speeds, indicating escape behaviors. All these changes in the urchin’s behavior decrease feeding opportunities and therefore reduce the grazing pressure on algal biomass. Furthermore, starved urchins seemed to be more insensitive and indifferent to predation cues. The 100% stay response from starved urchins despite the presence of a dead urchin in the field experiment further reinforced our hypothesis. Our findings suggest that the urchin’s hunger state was a key determinant in its decision-making process and that their level of hunger may override behaviors associated with predator avoidance. This puts them at a disadvantage as starved urchins feed more boldly, exposing themselves further to the dangers of predation.

## 6. ACKNOWLEDGEMENTS

We thank Yukio Inoue, Koichi Osaki and Kenjiro Hinode for assistance in the collection of seaweed samples. We also thank Mikiya Hidaka and Takeshi Urae for assistance in the collection of sea urchins. This study was supported by the Grant-in-Aid for Scientific Research (C-#40508321) from the Japan Society for the Promotion of Science (JSPS) and by the Japanese Ministry of Education, Culture, Sports, Science and Technology (MEXT).

## REFERENCES

Agatsuma Y, Nakata A, Matsuyama K (2000) Seasonal foraging activity of the sea urchin Strongylocentrotus nudus on coralline flats in Oshoro Bay in south-western Hokkaido, Japan. Fish Sci 66: 198–203.

Agatsuma Y (2013) Chapter 30: Hemicentrotus pulcherrimus, Pseudocentrotus depressus, and Heliocidaris crassispina. In Lawrence, J.M. (2013) (Ed.), Sea urchins: Biology and Ecology, Third Edition. Developments in Aquaculture and Fisheries Science, vol 38, Elsevier, United Kingdom, p 461–472.

Blamey LK, Plaganyi EE, Branch GM (2013) Modeling a regime shift in a kelp forest ecosystem caused by a lobster range expansion. Bull Mar Sci 89(1): 347–375.

Brockington S, Clarke A (2001) The relative influence of temperature and food on the metabolism of a marine invertebrate. J Exp Mar Biol Ecol 258: 87–99.

Clark CW (1994) Antipredator behavior and the asset-protection principle. Behavioral Ecology 5(2): 159–170.

Bonaviri C, Ferenandez TV, Fanelli G, Badalamenti F, Gianguzza P (2011) Leading role of the sea urchin Arbacia lixula in maintaining the barren state in southwestern Mediterranean. Mar Biol 158: 2505–2513.

Bürkner PC (2017) brms: An R package for Bayesian multilevel models using stan. J Stat Softw 80(1): doi: 10.18637/jss.v080.i01.

Campbell AC, Coppard S, D’Abreo C, Tudor-Thomas R (2001) Escape and aggregation responses of three echinoderms to conspecific stimuli. Biol Bull 201: 175–185.

Diaz de Vivar ME, Zarate EV, Rubilar T, Epherra L, Avaro MG, Sewell MA (2019) Lipid and fatty acid profiles of gametes and spawned gonads of Arbacia dufresnii (Echinodermata: Echinoidea): Sexual differences in the lipids of nutritive phagocytes. Mar Biol 166(96): doi.org/10.1007/s00227-019-3544-y.

Estes JA, Palmisano JF (1974) Sea otters: Their role in structuring nearshore communities. Science 185(4156): 1058–1060.

Feng WP, Nakabayashi N, Narita K, Inomata E, Aoki MN, Agatsuma Y (2019) Reproduction and population structure of the sea urchin Heliocidaris crassispina in its newly extended range: The Oga Peninsula in the Sea of Japan, northeastern Japan. PLoS ONE 14(1): doi.org/10.1371/journal.pone.0209858.

FFmpeg Developers (2018) FFmpeg version 4.1.1. [software]. Available from http://www.ffmpeg.org/.

Filbee-Dexter K, Scheibling RE (2014) Sea urchin barrens as alternative stable states of collapsed kelp ecosystems. Mar Ecol Prog Ser 495: 1–25.

Filbee-Dexter K and Wernberg T (2018) Rise of turfs: A new battlefront for globally declining kelp forests. BioSci, 68(2): 64–76.

Finelli CM, Pentcheff ND, Zimmer-Faust RK, Wethey DS (1999) Odor transport in turbulent flow: constraints on animal navigation. Limnol Oceanogr 44(4): 1056–1071.

Fletcher D, Mackenzie D, Villouta E (2005) Modelling skewed data with many zeros: A simple approach combining ordinary and logistic regression. Environ Ecol Stat 12: 45–54.

Flukes EB, Johnson CR, Ling SD (2012) Forming sea urchin barrens from the inside out: an alternative pattern of overgrazing. Mar Ecol Prog Ser 464: 179–194.

Fraser DF, Huntingford FA (1986) Feeding and avoiding predation hazard: the behavioral response of the prey. Ethology 73: 56–68.

Fujita D (2010) Current status and problems of isoyake in Japan. Bull Fish Res Agency 32: 33–42.

Fujita D (2015) Toward the revision of isoyake taisaku guideline. Fish Eng 51(3): 259–262.

Goodrich B, Gabry J, Ali I, Brilleman S (2018) rstanarm: Bayesian applied regression modeling via Stan. http://mc-stan.org/.

Hagen NT, Andersen A, Stabell OB (2002) Alarm responses of the green sea urchin, Strongylocentrotus droebachiensis, induced by chemically labelled durophagous predators and simulated acts of predation. Mar Biol 140: 365–374.

Hairston NG, Smith FE, Slobodkin LB (1960) Community structure, population control and competition. Am Nat 94(879): 421–425.

Halaj J, Wise DH (2001) Terrestrial and trophic cascades: How much do they trickle? Am Nat 157(3): 262–281.

Haraguchi H, Sekida S (2008) Recent changes in the distribution of Sargassum species in Kochi, Japan. Kuroshio Sci 2(1): 41–46.

Harding APC, Scheibling RE (2015) Feed or flee: Effect of a predation risk cue on sea urchin foraging activity. J Exp Mar Biol Ecol 466: 59–69.

Hernandez-Carmona G, Garcia O, Robledo D, Foster M (2000) Restoration techniques for Macrocystis pyrifera (Phaeophyceae) populations at the southern limit of their distribution in Mexico. Bot Mar 43: 273–284.

Holbrook SJ, Carr MH, Schmitt RJ, Coyer JA (1990) Effect of giant kelp on local abundance of reef fishes: The importance of ontogenetic resource requirements. Bull Mar Sci 47(1): 104–114.

Johnson CR, Chabot RH, Marzloff MP, Wotherspoon S (2016) Knowing when (not) to attempt ecological restoration. Restor Ecol 25(1): 140–147.

Kaehler S, Kennish R (1996) Summer and winter comparisons in the nutritional value of the marine macroalgae from Hong Kong. Bot Mar 39: 11–17.

Kamimura Y, Shoji J (2009) Seasonal changes in the fish assemblage in a mixed vegetation area of seagrass and macroalgae in the central Seto Inland Sea. Aquacult Sci 57(2): 233–241.

Kaneko I, Ikeda Y, Ozaki H (1981) Biometrical relationships between body weight and organ weights in freshly sampled and starved sea urchins. Bull Jap Soc Fish 47: 539–537.

Kassahun W, Neyens T, Molenberghs G, Faes C, Verbeke G (2014) Marginalized multilevel hurdle and zero-inflated models for overdispersed and correlated count data with excess zeros. Statist Med, 33: 4402–4419. DOI: 10.1002/sim.6237.

Kiyomoto S, Tagawa M, Nakamura Y, Horii T, Watanabe S, Tozawa T, Yatsuya K, Yoshimura T, Tamaki, A (2013) Decrease of abalone resources with disappearance of macroalgal beds around the Ojika Islands, Nagasaki, Southwestern Japan. J Shellfish Res 32(1): 51–58.

Kriegisch N, Reeves SE, Flukes EB, Johnson CR, Ling SD (2019) Drift-kelp suppresses foraging movement of overgrazing urchins. Oecol. https://doi.org/10.1007/s00442-019-04445-6

Kuwahara H, Hashimoto O, Sato A, Fujita D (2010) Introduction of isoyake recovery guideline (Fisheries Agency, Japan). Bull Fish Res Agency 32: 51–60.

Langlois TJ, Radford BT, Van Niel KP, Meeuwig JJ, Pearce AF, Rousseaux CSG, Kendrick GA, Harvey ES (2011) Consistent abundance distributions of marine fishes in an old, climatically buffered, infertile seascape. Glob Ecol Biogeogr 21(9): 886–897. http://dx.doi.org/10.1111/j.1466-8238.2011.00734.x.

Lauzon-Guay JS, Scheibling RE (2007) Behaviour of sea urchin Strongylocentrotus droebachiensis grazing fronts: food mediated aggregation and density-dependent facilitation. Mar Ecol Prog Ser 329: 191–204.

Lawrence JM (1970) The effects of starvation on the lipid and carbohydrate levels of the gut of the tropical sea urchin Echinometra mathaei (de Blainville). Pac Sci, 24: 487–489.

Layton C, Coleman MA, Marzinelli EM, Steinberg PD, Swearer SE, Verges A, Wernberg T, Johnson CR (2020) Kelp forest restoration in Australia. Front Mar Sci, 7:74. Doi: 10.3389/fmars.2020.00074.

Lewin WC, Freyhof J, Huckstorf V, Mehner T, Wolter C (2010) When no catches matter: Coping with zeros in environmental assessments. Ecol Ind, 10: 572–583. Doi: 10.1016/j.ecolind.2009.09.006.

Ling SD, Johnson CR (2009) Population dynamics of an economically important range-extender: kelp beds versus sea urchin barrens. Mar Ecol Prog Ser 374: 113–125.

Ling SD, Ibbott S, Sanderson JC (2010) Recovery of canopy-forming macroalgae following removal of the enigmatic grazing sea urchin Heliocidaris erythrogramma. J Exp Mar Biol Ecol 395: 135–146.

Ling SD, Scheibling RE, Rassweiler A, Johnson CR, Shears N, Connell SD, Salomon AK, Norderhaug KM, Perez-Matus A, Hernandez JC, Clemente S, Blamey LK, Hereu B, Ballesteros E, Sala E, Garrabou J, Cebrian E, Zabala M, Fujita D, Johnson LE (2015) Global regime shift dynamics of catastrophic sea urchin overgrazing. Phil Trans B 370: 20130269. http://dx.doi.org/10.1098/rstb.2013.0269.

Lowry LF, Pearse JS (1973) Abalones and sea urchins in an area in habited by sea otters. Mar Biol 23: 213–219.

Luttberg B, Rowe L, Mangel M (2003) Prey state and experimental design affect relative size of trait- and density-mediated indirect effects. Ecology 84(5): 1140–1150.

Lynam CP, Llope M, Möllmann C, Helaouët P, Bayliss-Brown GA, Stenseth NC (2017) Interaction between top-down and bottom-up control in marine food webs. Proc Nat Acad Sci 114(8): 1952–1957.

Magurran AE, Girling SL (1986) Predator model recognition and response habituation in shoaling minnows. Anim Behav 34: 510–518.

Mangialajo L, Chiantore M, Cattaneo-Vietti R (2008) Loss of fucoid algae along a gradient of urbanisation, and structure of benthic assemblages. Mar Ecol Prog Ser 358: 63–74.

Martinez-Pita I, Garcia FJ, Pita ML (2010) The effects of seasonality on gonad fatty acids of the sea urchins Paracentrotus lividus and Arbacia lixula (Echinodermata: Echinoidea). J Shellfish Res 29(2): 517–525.

Menge BA, Sutherland JP (1987) Community regulation: Variation in disturbance, competition, and predation in relation to environmental stress and recruitment. Am Nat 130(5): 730–757.

Morishita VM, Barreto RE (2011) Black sea urchins evaluate predation risk using chemical signals from a predator and injured con- and heterospecific prey. Mar Ecol Prog Ser 435: 173–181.

Nanri K, Nakajima Y, Yatzuya K, Kiyomoto S, Andou W, Yoshimura T (2011) Some approaches for the recovery from barren grounds in Shin-Mie, Nagasaki Prefecture. Fish Eng 48(1): 59–64.

Nielsen KJ, Navarrete SA (2004) Mesoscale regulation comes from the bottom-up: intertidal interactions between consumers and upwelling. Ecol Lett 7: 31–41.

Ogata R, Takagi K, Mishiku A, Fujita D (2016) A trial of restoration of a Sargassum forest by transplanting thalli with a suspended net in Uchiura Bay, Shizuoka Prefecture. Fish Eng 52(3): 177–184.

Okuda K (2008) Coastal environment and seaweed-bed ecology in Japan. Kuroshio Sci 2(1): 15–20.

Peacor SD, Werner EE (2001) The contribution of trait-mediated indirect effects to the net effects of a predator. Proc Nat Acad Sci, 98(7): 3904–3908.

Pearse JS, Hines AH (1987) Long-term population dynamics of sea urchins in a central California Kelp forest: rare recruitment and rapid decline. Mar Ecol Prog Ser, 39: 275–283.

Pedersen HG, Johnson CR (2008) Growth and age structure of sea urchins (Heliocidaris erythrogramma) in complex barrens and native macroalgal beds in eastern Tasmania. ICES J Mar Sci 65(1): 1–11.

Pessarrodona, A, Boada J, Pages JF, Arthur R, Alcoverro T (2019) Consumptive and non-consumptive effects of predators vary with the ontogeny of their prey. Ecol, 100(5): 302649.doi:10.1002/ecy.2649.

R Development Core Team 2019. A language and environment for statistical computing. R Foundation for Statistical Computing, Vienna, Austria. http://www.R-project.org.

Rocha F, Baiao LF, Moutinho S, Reis B, Oliveira A, Arenas F, Maia MRG, Fonseca AJM, Pintado M, Valente LMP (2019) The effect of sex, season and gametogenic cycle on gonad yield, biochemical composition and quality traits of Paracentrotus lividus along the North Atlantic coast of Portugal. Sci Rep 9(2994). doi.org/10.1038/s41598-019-39912-w.

Rodriguez-Barreras R, Cuevas E, Cabanillas-Teran N, Sabat AM (2015) Potential omnivory in the sea urchin Diadema antillarum? Reg Stud Mar Sci 2: 11–18.

Sala E, Zabala M (1996) Fish predation and the structure of the sea urchin Paracentrotus lividus populations in the NW Mediterranean. Mar Ecol Prog Ser 140: 71–81.

Scheibling RE, Anthony SX (2001) Feeding, growth and reproduction of sea urchins (Strongylocentrotus droebachiensis) on single and mixed diets of kelp (Laminaria sp.) and the invasive alga Codium fragile ssp. tomentosoides. Mar Biol 139: 139–146.

Schmitz OJ, Krivan V, Ovadia O (2004) Trophic cascades: The primacy of trait-mediated indirect interactions. Ecol Lett 7: 153–163.

Shurin JB, Borer ET, Seabloom EW, Anderson K, Blanchette CA, Broitman B, Cooper SD, Halpern BS (2002) A cross-ecosystem comparison of the strength of trophic cascades. Ecol Lett 5: 785–791.

Sievers D, Nebelsick JH (2018) Fish predation on a Mediterranean echinoid: Identification and preservation potential. Palaios 33: 47–54.

Smale DA, Kendrick GA, Wernberg T (2010) Assemblage turnover and taxonomic sufficiency of subtidal macroalgae at multiple spatial scales. J Exp Mar Biol Ecol 384: 76–86.

Smee DL, Weissberg MJ (2006) Clamming up: environmental forces diminish the perceptive ability of bivalve prey. Ecology 87(6): 1587–1598.

Spyksma AJP, Shears, NT, Taylor RB (2020) Injured conspecifics as an alarm cue for the sea urchin Evechinus chloroticus. Mar Ecol Prog Ser 641:135–144.

Stan Development Core Team (2019) RStan: The R interface to Stan. http://mc-stan.org/.

Steneck RS, Graham MH, Bourque BJ, Corbett D, Erlandson JM, Estes JA, Tegner, MJ (2002) Kelp forest ecosystems: Biodiversity, stability, resilience and future. Environ Conserv 29(4): 436–459.

Steneck RS, Leland A, McNaught DC, Vavrinec J (2013) Ecosystem flips, locks and feedbacks: The lasting effects of fisheries on Maine’s kelp forest ecosystem. Bull Mar Sci 89(1): 31–55.

Strong DR (1992) Are trophic cascades all wet? Differentiation and donor-control in speciose ecosystems. Ecology 73(3): 747–754.

Tegner MJ, Levin LA (1983) Spiny lobsters and sea urchins: Analysis of predator-prey interaction. J Exp Mar Biol Ecol 73: 125–150.

Terawaki T, Yoshikawa K, Yoshida G, Ushimura M, Iseki K (2003) Ecology and restoration techniques for Sargassum beds in the Seto Inland Sea, Japan. Mar Poll Bull 47: 198–201.

Tuya F, Boyra A, Sanchez-Jerez P, Barbera C, Haroun R (2004) Can one species determine the structure of the benthic community on a temperate rocky reef? The case of the long-spined sea-urchin Diadema antillarum (Echinodermata: Echinoidea) in the Eastern Atlantic. Hydrobiol, 519: 211–214.

Uki N, Sugiura M, Watanabe T (1986) Dietary value of seaweeds occurring on the Pacific Coast of Tohoku for growth of the abalone Haliotis discus hannai. Bull Jap Soc Sci Fish 52(2): 257–266.

Urriago JD, Wong JCY, Dumont CP, Qiu JW (2016) Reproduction of the short-spined sea urchin Heliocidaris crassispina (Echinodermata: Echinoidea) in Hong Kong with a subtropical climate. Reg Stud Mar 8: 445–453.

Vadas RL (1977) Preferential feeding: an optimization strategy in sea urchins. Ecol Monogr 47: 337–371.

Vanderklift MA, Kendrick GA (2005) Contrasting influence of sea urchins on attached and drift macroalgae. Mar Ecol Prog Ser 299: 101–110.

Verdura J, Sales M, Ballesteros E, Cefali ME, Cebrian E (2018) Restoration of a canopy-forming alga based on recruitment enhancement: Methods and long-term success assessment. Front Plant Sci, 9:1832. doi: 10.3389/fpls.2018.01832.

Verges A, Campbell AH, Wood G, Kajlich L, Eger AM, Cruz D, Langley M, Bolton D, Coleman MA, Turpin J, Crawford M, Coombes M, Camilleri A, Steinberg PD, Marzinelli EM (2020) Operation Crayweed: Ecological and sociocultural aspects of restoring Sydney’s underwater forests. Ecol Manag Restor, 21(2): 74–85. Doi: 10.1111/emr.12413.

Watanuki A, Yamamoto H (1990) Settlement of seaweeds on coastal structures. Hydrobiol 204/205: 275–280.

Watanuki A, Aota T, Otsuka E, Kawai T, Iwahashi Y, Kuwahara H, Fujita D (2010) Restoration of kelp beds on an urchin barren: Removal of sea urchins by citizen divers in southwestern Hokkaido. Bull Fish Res Agency 32: 83–87.

Webster DR, Weissberg MJ (2001) Chemosensory guidance cues in a turbulent chemical odor plume. Limnol Oceanogr 46(5): 1034–1047.

Westermeier R, Murua P, Patiño DJ, Muñoz L, Atero C, Müller DG (2013) Repopulation techniques for Macrocystis integrifolia (Phaeophyceae: Laminariales) in Atacama, Chile. J Appl Phycol 26(1): 511–518. DOI 10.1007/s10811-013-0069-5

Wright JT, Dworjanyn SA, Rogers CN, Steinberg PD, Williamson JE, Poore AGB (2005) Density-dependent sea urchin grazing: Differential removal of species, changes in community composition and alternative community states. Mar Ecol Prog Ser 298: 143–156.

Yamasaki M, Kiyomoto S (1993) Reproductive cycle of the sea urchin Anthocidaris crassispina from Hirado Island, Nagasaki Prefecture. Bull Seikai Nat Fish Res Inst 71: 33–40.

Yatsuya K, Nakahara H (2004a) Density, growth and reproduction of the sea urchin Anthocidaris crassispina (A. Agassiz) in two different adjacent habitats, the Sargassum area and Corallina area. Fish Sci 70: 233–240.

Yatsuya K, Nakahara H (2004b) Diet and stable isotope ratios of gut contents and gonad of the sea urchin Anthocidaris crassispina (A. Agassiz) in two different adjacent habitats, the Sargassum area and Corallina area. Fish Sci 70: 285–292.

Yoo SK, Hur SB, Ryu SY (1982) Growth and spawning of the sea urchin Anthocidaris crassispina (A. Agassiz). Bull Kor Fish Soc 15(4): 345–358.

Yoon JT, Sun SM, Chung G (2013) Sargassum bed restoration by transplanting of germlings grown under protective mesh cages. J Appl Phycol 26(1): 505–509. DOI 10.1007/s10811-013-0058-8.

Yotsui T, Maesako N (1993) Restoration experiments of Eisenia bicyclis beds on barren grounds at Tsushima Islands. Suisanzoshoku 41(1): 67–70.

Zimmer RK, Butman CA (2000) Chemical signaling processes in the marine environment. Biol Bull 198: 168–1187.

Zuur AF, Ieno EN, Walker NJ, Saveliev AA, Smith GM (2009) Zero-truncated and zero-inflated models for count data. In: Gail M, Krickeberg K, Samet JM, Tsiatis A, Wong W (eds) Mixed effects models and extensions in ecology with R. Statistics for Biology and Health. Pp. 261–293. Springer, New York, NY.

