## Supplementary for "Effects of dead conspecifics, hunger states, and seasons on the foraging behavior of the purple urchin *Heliocidaris crassispina*"

Supplementary Figures

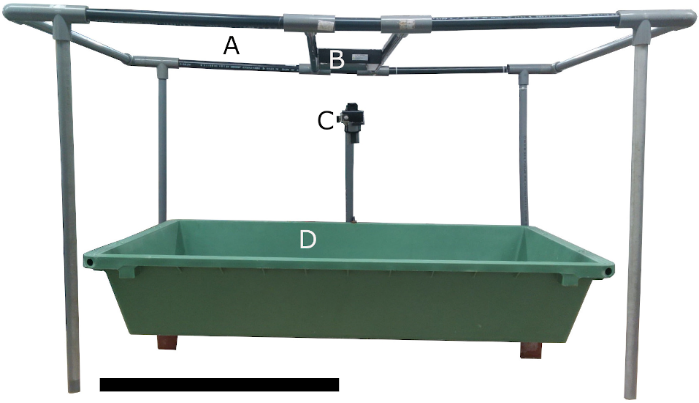

S1. The experiment 2 behavior experiment microcosm set-up. A) The PVC frame, B) is the red LED lamp mounted on the PVC frame, C) is the GoPro camera mounted over D) the microcosm tank. The black scale bar is 50 cm.

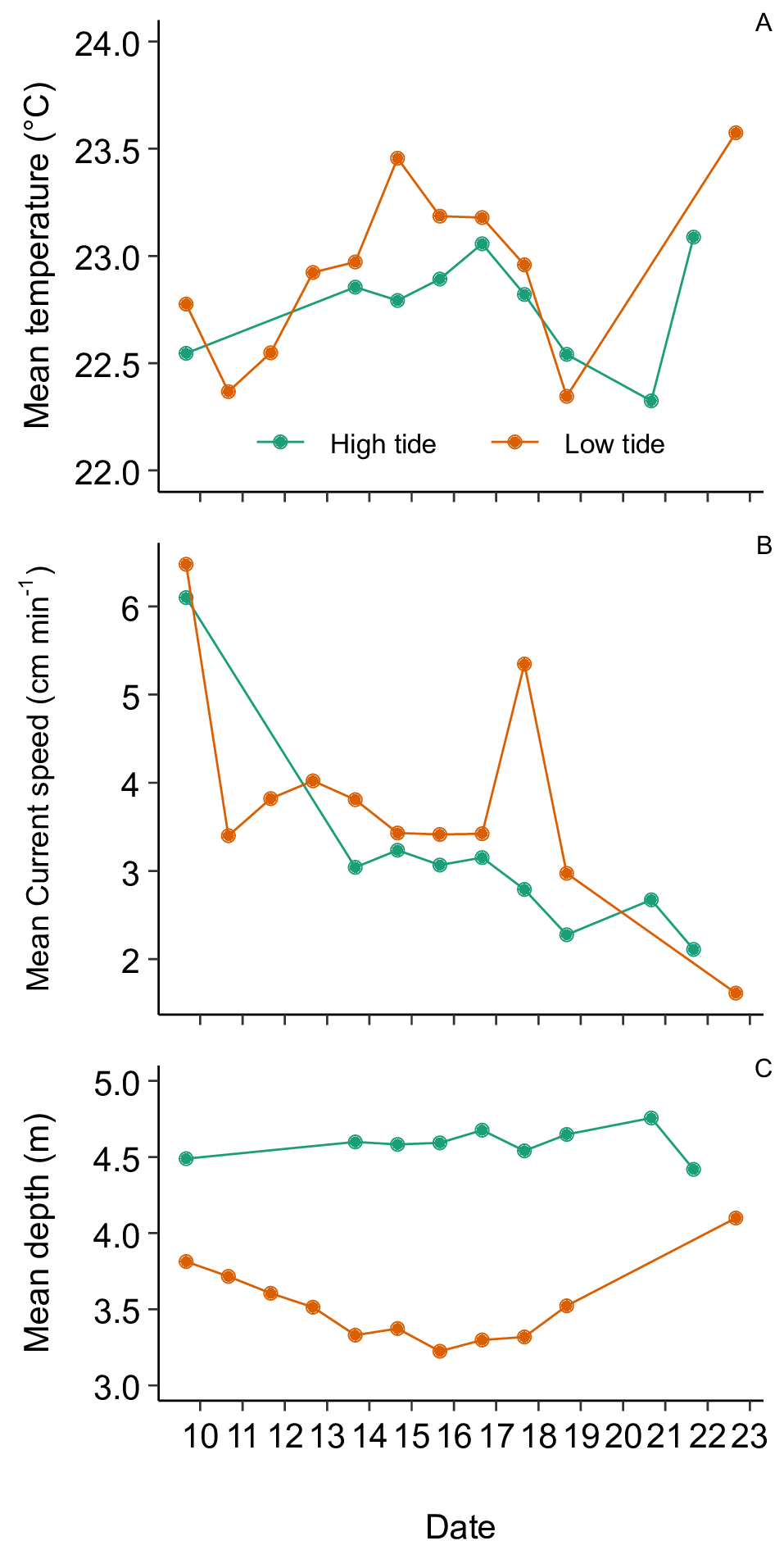

S2. The field experiment conditions in July 2019 at low and high tide. A) Shows the mean temperature conditions. B) Shows the mean current speed (cm sec^-1^), C) Shows mean depth. Points represent experiment times. The passing of tropical storm No. 5 (Danas) appears as a trough in the mean temperature and as an abrupt spike in mean current speed from the 18th to 19th of July 2019 (https://www.data.jma.go.jp/fcd/yoho/typhoon/route_map/bstv2019.html).

Supplementary table 1. Results of the Bayesian generalized linear models on light distribution of experiment 2 microcosm tank. The table shows the estimates, expected value and the lower and upper limits of the 95% highest density interval (HDI) of the expected value.

| Estimates | Expected value | 2.5% | 97.5% |
| --- | --- | --- | --- |
| **B1.) Light distribution (Lux)** | |  |  |
| Center | 3.1 | 2.8 | 3.3 |
| Middle | 2.1 | 2.0 | 2.3 |
| Outer | 0.5 | 0.3 | 0.7 |

Supplementary table 2. Mean and 95% highest density intervals (HDI) of the predictions of experiment 2 on sea urchin behavior counts.

|  |  | Summer-Autumn | | Winter-Spring | |
| --- | --- | --- | --- | --- | --- |
| Hunger state | Behavior | Mean | 95% HDI | Mean | 95% HDI |
| Control |  |  |  |  |  |
| Fed | Movement | 42.7 | 2-96 | 34.5 | 2-84 |
|  | Interaction | 21.8 | 2-52 | 22.3 | 1-52 |
|  | Outside | 48.2 | 3-119 | 49.6 | 3-116 |
| Starved | Movement | 45.5 | 3-116 | 45.6 | 2-107 |
|  | Interaction | 27.1 | 2-96 | 34.9 | 2-101 |
|  | Outside | 52.6 | 4-110 | 48.0 | 1-122 |
| Dead urchin |  |  |  |  |  |
| Fed | Movement | 29.3 | 2-67 | 44.9 | 2-107 |
|  | Interaction | 16.3 | 1-37 | 17.5 | 1-39 |
|  | Outside | 63.9 | 4-144 | 52.4 | 1-125 |
| Starved | Movement | 28.6 | 2-65 | 42.7 | 3-100 |
|  | Interaction | 22.9 | 1-53 | 22.8 | 1-56 |
|  | Outside | 62.0 | 1-140 | 56.7 | 3-137 |
| Algae |  |  |  |  |  |
| Fed | Movement | 23.3 | 1-54 | 32.8 | 3-73 |
|  | Interaction | 33.8 | 2-71 | 26.6 | 1-61 |
|  | Outside | 70.2 | 4-154 | 48.4 | 1-112 |
| Starved | Movement | 25.5 | 1-61 | 24.4 | 2-58 |
|  | Interaction | 66.7 | 1-161 | 66.0 | 1-163 |
|  | Outside | 67.1 | 3-173 | 58.9 | 3-153 |
| Dead urchin and algae | |  |  |  |  |
| Fed | Movement | 13.8 | 1-32 | 31.3 | 1-76 |
|  | Interaction | 12.9 | 1-32 | 8.3 | 1-21 |
|  | Outside | 76.3 | 3-183 | 60.6 | 3-140 |
| Starved | Movement | 21.4 | 1-50 | 15.0 | 1-35 |
|  | Interaction | 51.1 | 5-116 | 24.5 | 2-60 |
|  | Outside | 44.8 | 3-107 | 55.3 | 2-130 |

Supplementary table 3. Mean and 95% highest density intervals (HDI) of the predictions of experiment 2 on sea urchin behavior time (min).

|  |  | Summer-Autumn | | Winter-Spring | |
| --- | --- | --- | --- | --- | --- |
| Hunger state | Behavior | Mean | 95% HDI | Mean | 95% HDI |
| Control |  |  |  |  |  |
| Fed | Movement | 21.5 | 1.06-49.0 | 17.1 | 0.682-42.0 |
|  | Interaction | 10.6 | 0.570-24.2 | 10.9 | 0.165-26.7 |
|  | Outside | 22.9 | 1.23-50.9 | 25.1 | 0.958-61.9 |
| Starved | Movement | 23.2 | 1.49-59.0 | 23.2 | 0.815-54.6 |
|  | Interaction | 14.7 | 0.167-40.2 | 17.6 | 1.52-60.5 |
|  | Outside | 26.2 | 1.29-59.9 | 26.1 | 1.21-67.5 |
| Dead urchin |  |  |  |  |  |
| Fed | Movement | 15.0 | 0.994-35.1 | 22.6 | 0.848-52.9 |
|  | Interaction | 7.9 | 0.579-19.3 | 9.6 | 0.888-23.2 |
|  | Outside | 32.1 | 1.61-76.2 | 25.1 | 0.933-58.8 |
| Starved | Movement | 14.5 | 1.06-35.2 | 19.9 | 1.08-47.1 |
|  | Interaction | 10.6 | 0.400-27.7 | 12.2 | 0.271-33.6 |
|  | Outside | 30.8 | 1.42-74.2 | 28.9 | 2.09-77.4 |
| Algae |  |  |  |  |  |
| Fed | Movement | 11.8 | 0.738-27.6 | 17.0 | 0.844-38.6 |
|  | Interaction | 17.7 | 0.988-40.5 | 13.6 | 1.24-33.3 |
|  | Outside | 34.7 | 1.27-79.3 | 23.0 | 1.19-54.5 |
| Starved | Movement | 31.1 | 0.484-31.3 | 12.3 | 0.416-30.2 |
|  | Interaction | 33.4 | 1.61-82.1 | 33.2 | 0.982-81.1 |
|  | Outside | 31.1 | 2.54-82.1 | 29.1 | 0.772-75.7 |
| Dead urchin and algae | |  |  |  |  |
| Fed | Movement | 6.9 | 0.276-15.7 | 16.2 | 0.928-38.1 |
|  | Interaction | 6.5 | 0.180-14.7 | 4.3 | 0.180-10.6 |
|  | Outside | 38.8 | 2.19-91.2 | 30.6 | 0.645-73.5 |
| Starved | Movement | 10.8 | 0.568-25.3 | 7.3 | 0.366-17.5 |
|  | Interaction | 25.3 | 0.929-59.2 | 12.9 | 0.552-31.5 |
|  | Outside | 22.7 | 0.354-58.0 | 28.2 | 1.11-68.5 |

Supplementary table 4. Mean and 95% highest density intervals (HDI) of the predictions of experiment 2 on sea urchin behavior speed (cm sec ^-1^).

|  |  | Summer-Autumn | | Winter-Spring | |
| --- | --- | --- | --- | --- | --- |
| Hunger state | Behavior | Mean | 95% HDI | Mean | 95% HDI |
| Control |  |  |  |  |  |
| Fed | Movement | 7.22 | 0.106-19.8 | 12.5 | 0.094-35.6 |
|  | Interaction | 13.3 | 0.065-35.3 | 17.5 | 0.089-53.3 |
|  | Outside | 8.3 | 0.049-23.4 | 14.3 | 0.051-37.9 |
| Starved | Movement | 10.6 | 0.006-31.8 | 7.7 | 0.038-23.7 |
|  | Interaction | 11.1 | 0.082-33.7 | 8.5 | 0.138-23.3 |
|  | Outside | 4.2 | 0.063-14.4 | 3.5 | 0.028-11.2 |
| Dead urchin |  |  |  |  |  |
| Fed | Movement | 13.1 | 0.095-36.3 | 9.9 | 0.065-27.6 |
|  | Interaction | 21.9 | 0.098-62.2 | 16.9 | 0.092-48.4 |
|  | Outside | 6.7 | 0.065-18.5 | 9.5 | 0.076-28.5 |
| Starved | Movement | 18.1 | 0.416-50.8 | 11.2 | 0.087-32.2 |
|  | Interaction | 17.4 | 0.138-49.8 | 23.5 | 0.156-69.3 |
|  | Outside | 4.7 | 0.057-13.0 | 9.1 | 0.091-27.0 |
| Algae |  |  |  |  |  |
| Fed | Movement | 13.4 | 0.019-33.9 | 12.7 | 0.402-36.1 |
|  | Interaction | 16.6 | 0.243-46.5 | 12.5 | 0.164-34.3 |
|  | Outside | 13.4 | 0.026-37.2 | 12.7 | 0.072-33.8 |
| Starved | Movement | 8.4 | 0.026-24.5 | 8.6 | 0.017-24.0 |
|  | Interaction | 7.9 | 0.030-21.7 | 6.8 | 0.073-18.7 |
|  | Outside | 8.6 | 0.044-26.3 | 6.2 | 0.037-20.0 |
| Dead urchin and algae | |  |  |  |  |
| Fed | Movement | 20.1 | 0.081-57.0 | 15.0 | 0.007-38.6 |
|  | Interaction | 24.0 | 0.141-65.9 | 21.1 | 0.375-64.4 |
|  | Outside | 13.4 | 0.052-38.5 | 9.1 | 0.063-26.6 |
| Starved | Movement | 6.1 | 0.041-17.5 | 10.4 | 0.027-30.7 |
|  | Interaction | 8.3 | 0.043-23.0 | 21.4 | 0.302-60.9 |
|  | Outside | 22.8 | 0.065-70.3 | 14.7 | 0.123-42.8 |

Supplementary table 5. Mean and 95% highest density intervals (HDI) of the expected values on the probability of the behavior achieving zero counts.

|  |  | Summer-Autumn | | Winter-Spring | |
| --- | --- | --- | --- | --- | --- |
| Hunger state | Behavior | Mean | 95% HDI | Mean | 95% HDI |
| Control |  |  |  |  |  |
| Fed | Movement | 0.437 | 0.257-0.613 | 0.553 | 0.321-0.794 |
|  | Interaction | 0.752 | 0.624-0.876 | 0.818 | 0.670-0.939 |
|  | Outside | 0.593 | 0.414-0.774 | 0.635 | 0.399-0.853 |
| Starved | Movement | 0.738 | 0.517-0.925 | 0.570 | 0.304-0.817 |
|  | Interaction | 0.915 | 0.822-0.982 | 0.930 | 0.836-0.996 |
|  | Outside | 0.867 | 0.723-0.983 | 0.741 | 0.516-0.947 |
| Dead urchin |  |  |  |  |  |
| Fed | Movement | 0.180 | 0.034-0.344 | 0.371 | 0.147-0.614 |
|  | Interaction | 0.635 | 0.449-0.830 | 0.694 | 0.474-0.888 |
|  | Outside | 0.252 | 0.070-0.456 | 0.521 | 0.270-0.758 |
| Starved | Movement | 0.264 | 0.065-0.484 | 0.375 | 0.118-0.627 |
|  | Interaction | 0.723 | 0.531-0.915 | 0.821 | 0.625-0.978 |
|  | Outside | 0.387 | 0.158-0.617 | 0.566 | 0.286-0.827 |
| Algae |  |  |  |  |  |
| Fed | Movement | 0.234 | 0.093-0.383 | 0.398 | 0.197-0.608 |
|  | Interaction | 0.508 | 0.341-0.670 | 0.589 | 0.381-0.783 |
|  | Outside | 0.344 | 0.185-0.512 | 0.481 | 0.284-0.726 |
| Starved | Movement | 0.345 | 0.142-0.589 | 0.382 | 0.137-0.638 |
|  | Interaction | 0.567 | 0.343-0.786 | 0.697 | 0.437-.900 |
|  | Outside | 0.810 | 0.620-0.961 | 0.697 | 0.448-0.922 |
| Dead urchin and algae | |  |  |  |  |
| Fed | Movement | 0.116 | 0.007-0.260 | 0.434 | 0.188-0.699 |
|  | Interaction | 0.550 | 0.329-0.774 | 0.651 | 0.429-0.882 |
|  | Outside | 0.145 | 0.0143-0.303 | 0.441 | 0.170-0.693 |
| Starved | Movement | 0.083 | 0.002-0.227 | 0.306 | 0.072-0.561 |
|  | Interaction | 0.336 | 0.109-0.565 | 0.575 | 0.311-0.849 |
|  | Outside | 0.487 | 0.230-0.753 | 0.526 | 0.254-0.797 |

Supplementary table 6. Mean and 95% highest density intervals (HDI) of the expected values on the probability of the behavior time (min) being zero.

|  |  | Summer-Autumn | | Winter-Spring | |
| --- | --- | --- | --- | --- | --- |
| Hunger state | Behavior | Mean | 95% HDI | Mean | 95% HDI |
| Control |  |  |  |  |  |
| Fed | Movement | 0.436 | 0.269-0.636 | 0.546 | 0.310-0.788 |
|  | Interaction | 0.756 | 0.624-0.889 | 0.816 | 0.659-0.945 |
|  | Outside | 0.589 | 0.412-0.783 | 0.637 | 0.405-0.850 |
| Starved | Movement | 0.731 | 0.526-0.931 | 0.574 | 0.317-0.822 |
|  | Interaction | 0.915 | 0.826-0.989 | 0.931 | 0.835-0.995 |
|  | Outside | 0.867 | 0.723-0.987 | 0.746 | 0.509-0.947 |
| Dead urchin |  |  |  |  |  |
| Fed | Movement | 0.178 | 0.032-0.345 | 0.368 | 0.138-0.606 |
|  | Interaction | 0.638 | 0.447-0.823 | 0.695 | 0.483-0.905 |
|  | Outside | 0.254 | 0.068-0.442 | 0.524 | 0.266-0.774 |
| Starved | Movement | 0.263 | 0.064-0.483 | 0.377 | 0.121-0.634 |
|  | Interaction | 0.722 | 0.526-0.906 | 0.820 | 0.626-0.974 |
|  | Outside | 0.386 | 0.148-0.614 | 0.563 | 0.290-0.828 |
| Algae |  |  |  |  |  |
| Fed | Movement | 0.235 | 0.107-0.393 | 0.386 | 0.175-0.590 |
|  | Interaction | 0.503 | 0.344-0.659 | 0.589 | 0.401-0.795 |
|  | Outside | 0.345 | 0.187-0.507 | 0.483 | 0.266-0.699 |
| Starved | Movement | 0.340 | 0.119-0.563 | 0.386 | 0.130-0.645 |
|  | Interaction | 0.561 | 0.342-0.780 | 0.696 | 0.450-0.893 |
|  | Outside | 0.816 | 0.631-0.968 | 0.700 | 0.440-0.930 |
| Dead urchin and algae | |  |  |  |  |
| Fed | Movement | 0.118 | 0.007-0.262 | 0.437 | 0.173-0.677 |
|  | Interaction | 0.552 | 0.332-0.781 | 0.656 | 0.427-0.873 |
|  | Outside | 0.145 | 0.015-0.302 | 0.436 | 0.184-0.707 |
| Starved | Movement | 0.083 | 0.004-0.216 | 0.304 | 0.058-0.560 |
|  | Interaction | 0.341 | 0.124-0.580 | 0.577 | 0.323-0.836 |
|  | Outside | 0.493 | 0.248-0.772 | 0.522 | 0.249-0.800 |

Supplementary table 7. Mean and 95% highest density intervals (HDI) of the expected values on the probability of the behavior speeds (cm sec-1) being zero.

|  |  | Summer-Autumn | | Winter-Spring | |
| --- | --- | --- | --- | --- | --- |
| Hunger state | Behavior | Mean | 95% HDI | Mean | 95% HDI |
| Control |  |  |  |  |  |
| Fed | Movement | 0.501 | 0.326-0.704 | 0.547 | 0.299-0.776 |
|  | Interaction | 0.757 | 0.630-0.883 | 0.786 | 0.625-0.920 |
|  | Outside | 0.609 | 0.423-0.779 | 0.626 | 0.389-0.841 |
| Starved | Movement | 0.722 | 0.495-0.907 | 0.571 | 0.302-0.825 |
|  | Interaction | 0.893 | 0.792-0.980 | 0.915 | 0.805-0.992 |
|  | Outside | 0.864 | 0.718-0.984 | 0.744 | 0.506-0.940 |
| Dead urchin |  |  |  |  |  |
| Fed | Movement | 0.327 | 0.138-0.539 | 0.375 | 0.127-0.608 |
|  | Interaction | 0.701 | 0.524-0.879 | 0.745 | 0.548-0.915 |
|  | Outside | 0.310 | 0.110-0.524 | 0.537 | 0.280-0.784 |
| Starved | Movement | 0.299 | 0.102-0.534 | 0.369 | 0.129-0.640 |
|  | Interaction | 0.676 | 0.462-0.865 | 0.810 | 0.625-0.974 |
|  | Outside | 0.394 | 0.166-0.638 | 0.565 | 0.285-0.824 |
| Algae |  |  |  |  |  |
| Fed | Movement | 0.408 | 0.242-0.573 | 0.430 | 0.238-0.650 |
|  | Interaction | 0.551 | 0.387-0.701 | 0.585 | 0.393-0.775 |
|  | Outside | 0.345 | 0.184-0.509 | 0.480 | 0.256-0.690 |
| Starved | Movement | 0.441 | 0.212-0.691 | 0.387 | 0.134-0.650 |
|  | Interaction | 0.528 | 0.308-0.752 | 0.708 | 0.472-0.909 |
|  | Outside | 0.803 | 0.613-0.971 | 0.708 | 0.467-0.935 |
| Dead urchin and algae | |  |  |  |  |
| Fed | Movement | 0.594 | 0.331-0.819 | 0.559 | 0.298-0.820 |
|  | Interaction | 0.748 | 0.566-0.914 | 0.801 | 0.624-0.958 |
|  | Outside | 0.163 | 0.023-0.332 | 0.440 | 0.184-0.707 |
| Starved | Movement | 0.509 | 0.230-0.770 | 0.507 | 0.214-0.764 |
|  | Interaction | 0.413 | 0.172-0.652 | 0.658 | 0.405-0.904 |
|  | Outside | 0.494 | 0.226-0.748 | 0.519 | 0.228-0.773 |
